# Targeting high-risk multiple myeloma genotypes with optimized anti-CD70 CAR-T cells

**DOI:** 10.1101/2024.02.24.581875

**Authors:** Corynn Kasap, Adila Izgutdina, Bonell Patiño-Escobar, Amrik Kang, Nikhil Chilakapati, Naomi Akagi, Haley Johnson, Tasfia Rashid, Juwita Werner, Abhilash Barpanda, Huimin Geng, Yu-Hsiu T. Lin, Sham Rampersaud, Daniel Gil-Alós, Amin Sobh, Daphné Dupéré-Richer, Gianina Wicaksono, K.M. Kawehi Kelii, Radhika Dalal, Emilio Ramos, Anjanaa Vijayanarayanan, Fernando Salangsang, Paul Phojanakong, Juan Antonio Camara Serrano, Ons Zakraoui, Isa Tariq, Veronica Steri, Mala Shanmugam, Lawrence H. Boise, Tanja Kortemme, Elliot Stieglitz, Jonathan D. Licht, William J. Karlon, Benjamin G. Barwick, Arun P. Wiita

## Abstract

Despite the success of BCMA-targeting CAR-Ts in multiple myeloma, patients with high-risk cytogenetic features still relapse most quickly and are in urgent need of additional therapeutic options. Here, we identify CD70, widely recognized as a favorable immunotherapy target in other cancers, as a specifically upregulated cell surface antigen in high risk myeloma tumors. We use a structure-guided design to define a CD27-based anti-CD70 CAR-T design that outperforms all tested scFv-based CARs, leading to >80-fold improved CAR-T expansion in vivo. Epigenetic analysis via machine learning predicts key transcription factors and transcriptional networks driving CD70 upregulation in high risk myeloma. Dual-targeting CAR-Ts against either CD70 or BCMA demonstrate a potential strategy to avoid antigen escape-mediated resistance. Together, these findings support the promise of targeting CD70 with optimized CAR-Ts in myeloma as well as future clinical translation of this approach.

**One sentence summary:** Structure-optimized CD27-based CAR-T cells targeting CD70 are a promising therapeutic option for high-risk multiple myeloma patients who are most likely to relapse on current BCMA-targeting cellular therapies.

## Introduction

Multiple myeloma (MM) is a malignancy of plasma cells that remains without a definitive cure. Among recently-approved treatment modalities, the most promising have been bispecific antibodies and chimeric antigen receptor (CAR) T cells targeting the surface protein B-cell maturation antigen (BCMA) (*1*). However, even for highly effective BCMA-targeting CAR-Ts idecabtagene vicleucel (“ide-cel”) and ciltacabtagene-autoleucel (“cilta-cel”), it appears that most, if not all, patients will eventually relapse (*2*). Clinical studies for both CAR-T products in MM have found that one of the most prominent risk factors for early relapse is the presence of unfavorable or high-risk genomic features (*3, 4*). In parallel, other work has shown that patients relapsing after current BCMA CAR-Ts have poor outcomes with salvage regimens (*5*). Therefore, new therapeutic strategies for high-risk MM patients relapsing after BCMA CAR-T are urgently needed.

While not uniformly observed, mechanisms of resistance to BCMA CAR-T can include downregulation of surface BCMA and/or genetic deletion or mutation of the *TNFRSF17A* gene which encodes the BCMA protein (*6, 7*). In light of these findings, identification of alternate CAR-T targets has garnered significant interest in the MM field. The most clinically advanced in this space is the antigen GPRC5D (*8*). While GPRC5D-targeting CAR-Ts have shown highly promising initial responses, even in BCMA-relapsed patients, these responses have not yet proven to be durable (*9*). Additional CAR-T targets such as CD38, SLAMF7/CS-1, and CD138 have been proposed (*10*), but these antigens do not have high specificity for plasma cells, with moderate to high expression on numerous other normal cell types (*11*). These alternate targets therefore raise concerns about “on target, off tumor” toxicity when attacked with highly potent cellular therapies. There thus remains a desire to identify new CAR-T targets in MM with promising efficacy versus safety profiles.

One surface antigen that is known to be expressed on many tumors but with minimal expression on normal tissues is CD70 (*12, 13*). This surface marker is natively expressed only on limited subsets of T-cells, B-cells, NK-cells and dendritic cells after physiologic activation (*14*). Comprehensive analysis by immunohistochemistry reveals only rare CD70 positive cells in lymphoid structures, but no other normal human tissues (*15*). When expressed on normal hematopoietic cells, CD70 interacts in *trans* with the surface protein CD27 at the relevant cellular interface to serve as co-stimulatory signal (*14*). In contrast to this very limited expression on normal tissues, CD70 is known to be aberrantly expressed in several types of hematological malignancies, including acute myeloid leukemia (AML) (*16, 17*) and B-cell non-Hodgkin lymphoma (*18*), as well as many solid tumors, including renal cell carcinoma (*19*) and non-small cell lung cancer (*20*). Unifying mechanisms underlying CD70 upregulation across cancers remain unclear, although transcriptional programs enacted in the context of tyrosine kinase inhibitor resistance have been proposed in lung cancer (*20*) and chronic myelogenous leukemia (*21*). Preclinical studies in AML have suggested that CD70 is a promising CAR-T target for this malignancy (*22, 23*). A monoclonal antibody targeting CD70, cusatuzumab, has also shown a promising efficacy vs. safety profile in Phase I/II studies for AML (*16*). A phase I trial of a CAR-T cell targeting CD70 in renal cell carcinoma has also demonstrated an encouraging efficacy signal but with no sign of “on target, off tumor” toxicity (*24*). These findings validate CD70 as a promising cellular therapy target in cancer.

In MM, one prior study from 2008 used immunohistochemistry to evaluate CD70 expression across patient specimens using tissue microarrays, finding a heterogeneous expression pattern (*18*). While a CD70-targeting antibody drug conjugate showed efficacy versus MM cell lines (*18*), this strategy was not moved forward into clinical trials. In the intervening ~15 years, CD70 was largely forgotten as a potential MM immunotherapy target.

Here, in the BCMA CAR-T cell era, we seek to revisit CD70 as a potential therapeutic target. Transcriptional and flow cytometry analysis of MM patient tumors suggests that CD70 is particularly upregulated in several high-risk subtypes of the disease, in patients who appear most likely to relapse after BCMA CAR-T. Using a structure-guided design to optimize a CD27-based CAR-T versus CD70, we find that a truncated fragment of the CD27 extracellular domain has a greatly improved in vivo expansion and persistence as compared to standard antibody fragment (scFv) based CAR-T designs. Through epigenetic analysis, we further delineate key transcriptional regulators of CD70 in high-risk MM. Finally, we develop proof-of-principle dual-targeting CAR-Ts against both CD70 and BCMA, which could have utility in avoiding relapse due to loss or downregulation of either antigen. Collectively, these findings support pursuit of therapies directed against CD70 as a strategy for the growing numbers of MM patients relapsing after BCMA CAR-T.

## Results

### CD70 is upregulated in high-risk multiple myeloma

To identify new therapeutic targets in MM, we first sought to identify upregulated tumor cell surface targets in the broader context of relapsed/refractory MM. Toward this goal, we initially interrogated the Multiple Myeloma Research Foundation CoMMpass database (research.themmrf.org). We evaluated paired tumor RNA-seq data from patients (*n* = 50) both at initial diagnosis and then after relapse on standard induction therapy. It is important to note that none of the patients included in this analysis had received BCMA-targeting therapy, as CoMMpass data were collected before any such therapies were commercially available. However, in the absence of a similar publicly-available dataset after BCMA-directed therapy, we reasoned that targets upregulated in CoMMpass may provide clues to cell surface markers that are generally increased in resistant disease.

This analysis revealed 163 genes predicted to encode membrane-localized proteins (based on Uniprot annotation) that appeared significantly upregulated in relapsed MM plasma cells (**Fig. 1A**; **Supp. Table. 1**). Of these genes, we decided to focus on *CD70* (**Fig. 1A**) given its known promise as an immunotherapy target across other cancers, as discussed above. To confirm these findings through an orthogonal modality, we re-analyzed existing cell surface proteomics datasets of both MM cell lines and primary patient CD138+ tumor cells (*25, 26*). We indeed identified CD70 protein to be consistently present across MM cell lines, at proteomic intensities similar to that of AML cell lines analyzed in parallel (**Fig. S1A-B**). Surface proteomics revealed detectable CD70 peptides in 9 of 15 (60%) primary MM tumors (**Fig. S1A**).

**Figure 1.**
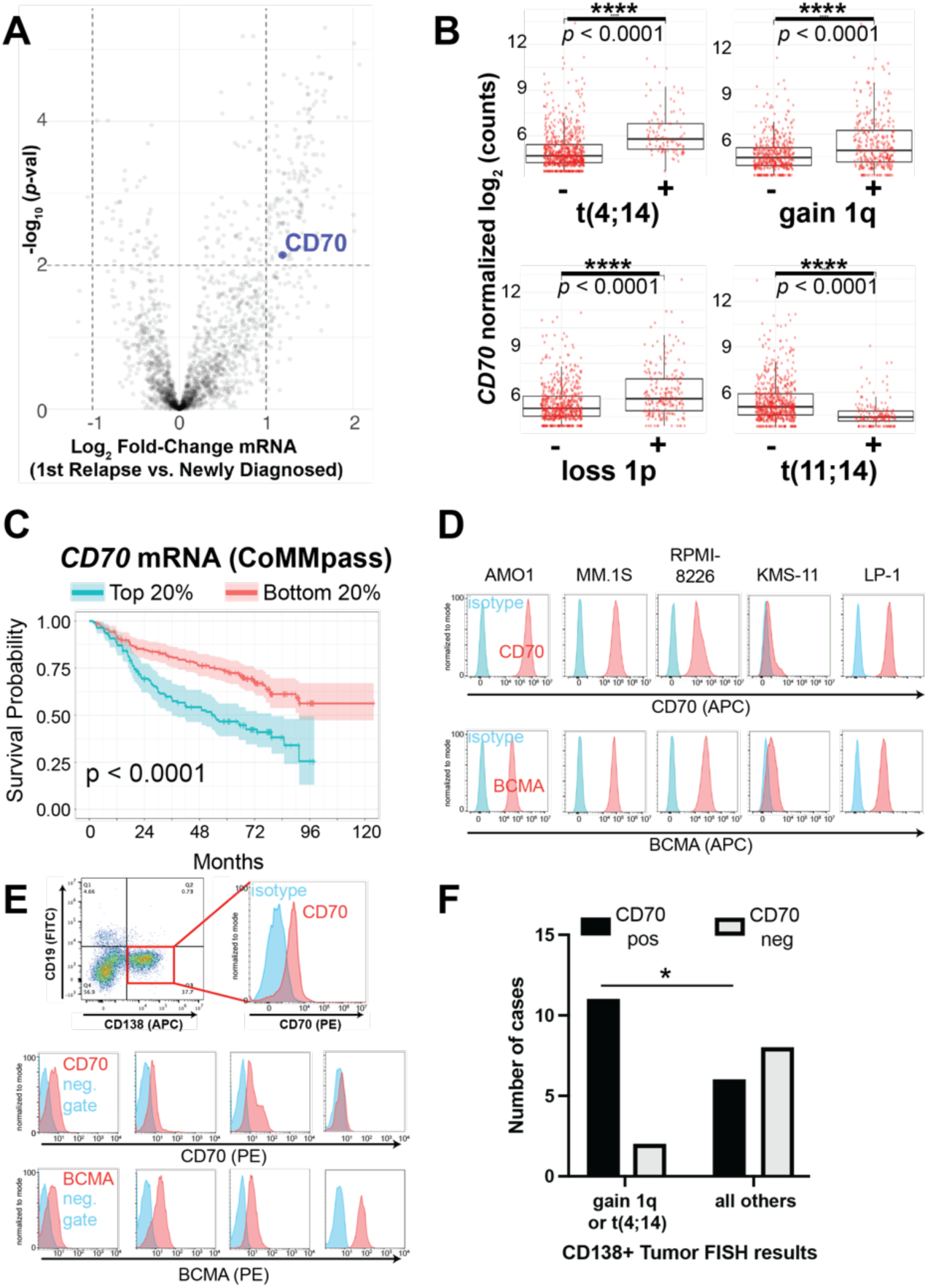
CD70 is an upregulated surface antigen in high-risk myeloma subtypes. **A.** Volcano plot of transcripts in the CoMMpass tumor database (*n* = 50 patients, paired relapse vs. diagnosis) filtered on those encoding plasma membrane proteins. Significance cutoff *p* < 0.01, log_2_FC > |1|. **B.** *CD70* expression in newly-diagnosed MM tumors in CoMMpass (*n* = 776; release IA19) as a function of genotype. *p*-value by two-tailed *t*-test. **C.** Overall survival in CoMMpass patients selected for the top and bottom quintile of CD70 expression. *p*-value by log-rank test. **D.** Flow cytometry on a panel of MM cell lines comparing CD70 and BCMA expression. Representative of *n* = 2 per line. **E.** Flow cytometry on fresh MM patient tumor samples selected as CD138+/CD19− in bone marrow aspirate (top example; isotype used as negative control) or selected as CD38+/CD45 dim/-/CD19− (bottom examples; fluorescence minus one (FMO) used as negative gate). Additional patient sample data shown in Fig. S2. 13 of 27 samples had both BCMA and CD70 measured. Each sample measured once due to sample input limitations. **F.** Fraction of patient tumor samples defined as positive for CD70 by flow cytometry (MFI >2x that of negative control) as a function of tumor genotype determined on diagnostic fluorescence in situ hybridization (FISH) per the clinical record. *p*-value by Fisher’s exact test. \**p*<0.05.

Deeper analysis of RNA-Seq data from a larger cohort of newly diagnosed patients from the CoMMpass data set (*n* = 776) showed that patients with high risk MM, as defined by Revised-International Staging System (R-ISS) criteria (*27*) at diagnosis, tended to have tumors with significantly increased *CD70* expression (*p* = 0.004 by *t* test, R-ISS 3 vs. R-ISS 1) (**Fig. S1C**). Further analysis based on well-characterized tumor genotypes showed that several high-risk MM karyotypic subtypes, specifically those with t(4;14), gain of Chr1q, and loss of Chr1p (*27, 28*), showed significantly upregulated *CD70* expression compared to all others in CoMMpass (**Fig. 1B**). In contrast, the standard risk karyotype of t(11;14) showed significantly decreased *CD70* expression (**Fig. 1B**). Additional analyses of CoMMpass data in the context of 7 transcriptionally-defined MM subtypes (*29*) confirmed that *CD70* was highest in the “MMSET” (MS) subtype, defined by t(4;14) tumors and associated with elevated *NSD2* expression (**Fig. S1D**). Cross correlation of *CD70* with all expressed genes in the CoMMpass cohort followed by subsequent Gene Set Enrichment Analysis identified a strong enrichment of gene signatures associated with cell cycle progression, *myc*-regulated networks, and genes localized to Chr1q, with de-enrichment of genes localized to Chr13p (**Fig. S1E**). Increased *CD70* expression was broadly associated with decreased overall survival in the CoMMpass cohort (**Fig. 1C**). These findings suggest that several groups of high-risk MM patients could benefit from CD70-targeting immunotherapies.

To further evaluate CD70 surface protein expression in MM we turned to flow cytometry. Analysis of several MM cell lines showed a range of CD70 expression (**Fig. 1D**), with similar number of CD70 molecules per cell compared to BCMA (**Fig. S2A**). Analysis of primary MM patient bone marrow aspirate further confirmed CD70 expression on 17 of 27 (63.0%) CD138+/CD38+/CD19− plasma cell specimens (**Fig. 1E**, **Fig. S2B-C**), based on a metric of mean fluorescence intensity (MFI) ratio >2 versus negative control. For the subset of samples with available BCMA staining, by this same metric we found 11 of 13 (84.6%) as BCMA positive (**Fig. 1E**, **Fig. S2B**). Overall, on primary samples BCMA intensity was moderately higher than CD70 (**Fig. S2D**); for both antigens, staining intensity was lower on primary samples than cell lines. In concordance with our transcriptomic analysis, comparison to available FISH data showed increased fraction of CD70 positivity for samples with gain of Chr1q (9 of 11 positive) or t(4;14) (2 of 2 positive) compared to other genotypes (**Fig. 1F**). Taken together, these results suggest that CD70 is a viable surface immunotherapy target for a majority of MM patients, and particularly those with the noted high-risk tumor genotypes.

### CRD1 of CD27 is required in an anti-CD70 CAR construct

After validating CD70 expression on primary MM, we next sought to develop a strategy to target this antigen therapeutically. Given the success of other CAR-T cells in MM, as well as the potential for CAR-Ts targeting CD70 in other cancers (*22, 23, 30, 31*), we aimed to test several CD70 CAR-T variants to identify one with optimal performance in MM models. To do this, we evaluated a broad range of 17 potential CAR-T designs (**Fig. 2A**). Using antibody fragments published in the patent literature as well as the sequence of clinically-tested monoclonal antibody cusatuzumab (*16*), we first developed 10 different single chain variable fragment (scFv) based CAR designs (discussed in more detail below). Such scFv’s are the most standard strategy for CAR-T design in the field, and are used in 5 of 6 commercially-available CAR-T products (*32*).

**Figure 2.**
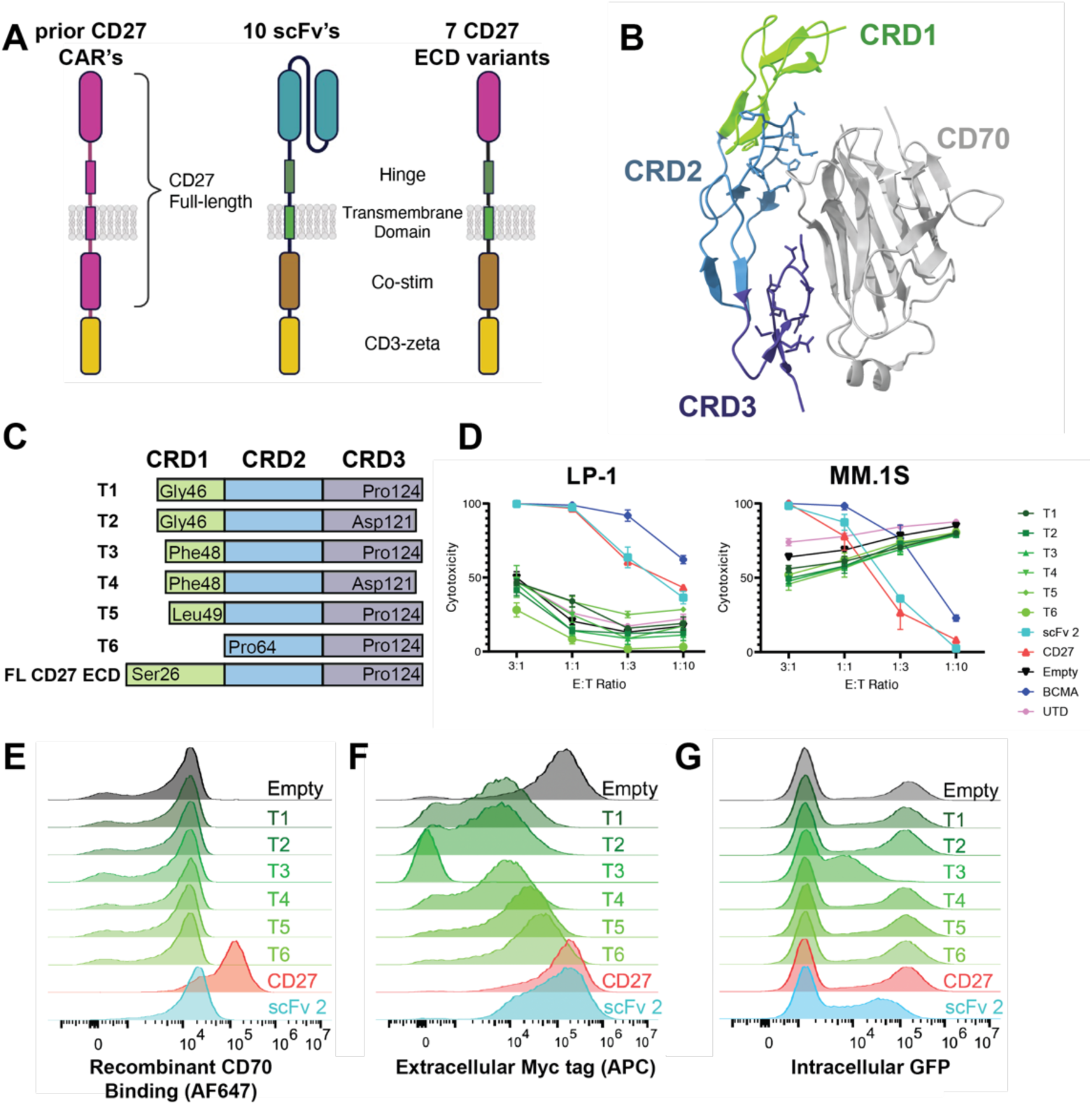
CRD1 of CD27 is required to generate functional anti-CD70 CAR-Ts. **A.** Illustration of various anti-CD70 CAR-T designs. Several prior CD70 CARs have included a full-length CD27 sequence fused to CD3z (left). Here we probe 10 different scFv-based CARs (sequences in Table S3) and several CD27 extracellular domain (ECD) variants fused to a CD28-based CAR backbone. **B.** Crystal structure of CD27:CD70 complex (PDB: 7XK0) illustrating domains CRD1-3 (“cysteine rich domains”) of CD27 interacting with the CD70 ECD. **C.** CD27 ECD truncation designs (T1-T6) used in tested CAR constructs. “FL” = full-length. **D.** In vitro cytotoxicity assays comparing tumor lysis of different CD27 CAR designs compared to positive control of anti-BCMA CAR, “scFv 2” anti-CD70 CAR, and negative controls of “empty” CAR (no antigen binding domain) and “UTD” (untransduced T-cells). 18 hrs, lysis measured by luciferase at the indicated effector:tumor (E:T) ratios. *n* = 4 technical replicates, representative of 2 biological replicates. Note the elevated lysis of MM.1S cells by truncated constructs is an artifact of increased MM.1S growth in the normalized “tumor only” wells, as evidenced by no dose-response at higher E:T. **E.** Flow cytometry on lentivirally transduced donor T-cells reflecting binding to FITC-conjugated recombinant CD70. **F.** Flow cytometry of *myc* tag present at the N-terminus (extracellular) of all CAR constructs. **G.** Flow cytometry of intracellular GFP present on CAR construct. E-G: representative of 2 biological replicates. “CD27” = FL ECD CD27 construct.

In parallel, we investigated the potential for using the extracellular domain of CD27 as the antigen recognition element for CD70. CD27-based anti-CD70 CAR-T designs had previously been explored in prior studies and shown potential for efficacy (*23, 30, 33*). In these studies, the CAR typically consists of a full length CD27 sequence (extracellular domain, transmembrane, and intracellular domains all derived from native CD27 (**Fig. 2A**)) fused to a CD3z domain. Here, we instead took advantage of the recently-published crystal structure of the CD27:CD70 hexameric complex (PDB: 7KX0) (*34*). We aimed to define a minimal fragment of the CD27 extracellular domain (ECD) required to engage CD70 in a functional CAR, when otherwise fused to a more typical CAR backbone (IgG4-based spacer, CD28 transmembrane, CD28 costimulatory domain, CD3z (**Fig. 2A**)). In each construct, GFP is included after a T2A element to monitor transduction efficiency.

Based on the CD27:CD70 crystal structure, we noted that the majority of the CD27 residues contacting CD70 were present in cysteine rich domain (CRD) 2 and 3, with a minimal contribution of CRD1 (**Fig. 2B**). Therefore, we designed a series of 6 truncations of the CD27 extracellular domain ranging from a full deletion of CRD1 to a deletion of only the N-terminal 20 amino acids of CRD1 (**Fig. 2C**). We lentivirally transduced donor T-cells and used a Cas9 ribonucleoprotein (RNP)-based strategy to knock out *CD70* in T-cells to prevent fratricide during manufacturing (**Fig. S3A**), as shown by others to improve anti-CD70 CAR-T performance (*35*). Surprisingly, *in vitro* cytotoxicity assays against both MM.1S and LP-1 MM cell lines found potent anti-tumor activity for the full-length CD27 ECD-based CAR, but no detectable activity for any of the CRD1 truncations (**Fig. 2D**). To further investigate this unexpected finding, we probed whether these different CAR constructs could be stained by fluorescently-labeled recombinant CD70 at the T-cell surface. In line with the *in vitro* cytotoxicity assays, we found that while full length CD27 ECD-based CAR engaged recombinant CD70, none of the CRD1 truncation constructs could do so (**Fig. 2E**). An anti-CD27 antibody also did not stain any of the truncation constructs (**Fig. S3B**). These findings raised the question of whether the truncations were translocated to the T-cell surface at all. Using an extracellular MYC tag on the CAR construct, we confirmed that all but one of these truncation CAR constructs was indeed present at the T-cell surface, albeit at lower levels than the full-length CD27 ECD (**Fig. 2F**). Relative intracellular GFP levels were similar to extracellular MYC staining (**Fig. 2G**). Together, these findings suggest decreased translation and/or increased degradation of the CRD1-truncated CAR constructs.

In line with our experimental results, structural modeling using Rosetta (*36*) suggested that elimination of some or all of CRD1 will variably decrease affinity for CD70 (**Fig. S3C-D**). While our work was in progress, Maus and colleagues published findings confirming that the full length CD27 ECD fused to a 4-1BB-based CAR construct could generate potent anti-CD70 CAR-Ts for use in AML (*22*). Notably, a patent filing extending from that work suggested that only a 12 amino acid fragment within CRD2 could serve as a minimal fragment to engage CD70 in a CAR (*37*). However, our work here delineates that the CRD2 sequence alone is insufficient to generate a functional CAR; CRD1 of CD27 is required.

### scFv- and CD27-based CAR-T designs have similar anti-myeloma activity in vitro

We next moved to compare the efficacy of the scFv-based constructs to the full-length CD27 ECD-based CAR-T (hereafter referred to as “CD27-based”). We tested our 10 scFv-based CARs (sequences in **Table S4**) in an initial screen versus the AMO-1 cell line (**Fig. S4A**). We further chose a subset of the most potent scFv-based CARs to evaluate anti-tumor effects *in vitro* (noted as scFv-2, -3, and -4), in comparison to CD27-based CAR-T as well as anti-BCMA CAR-T positive control (BCMA binder equivalent sequence to cilta-cel). Evaluation of these CAR-Ts demonstrated anti-tumor activity versus several MM cell lines (**Fig. 3A**) with varying CD70 antigen density (**Fig. 1D**). Encouragingly, activity of all CD70-targeting CAR-Ts was generally comparable to anti-BCMA CAR-T.

**Figure 3.**
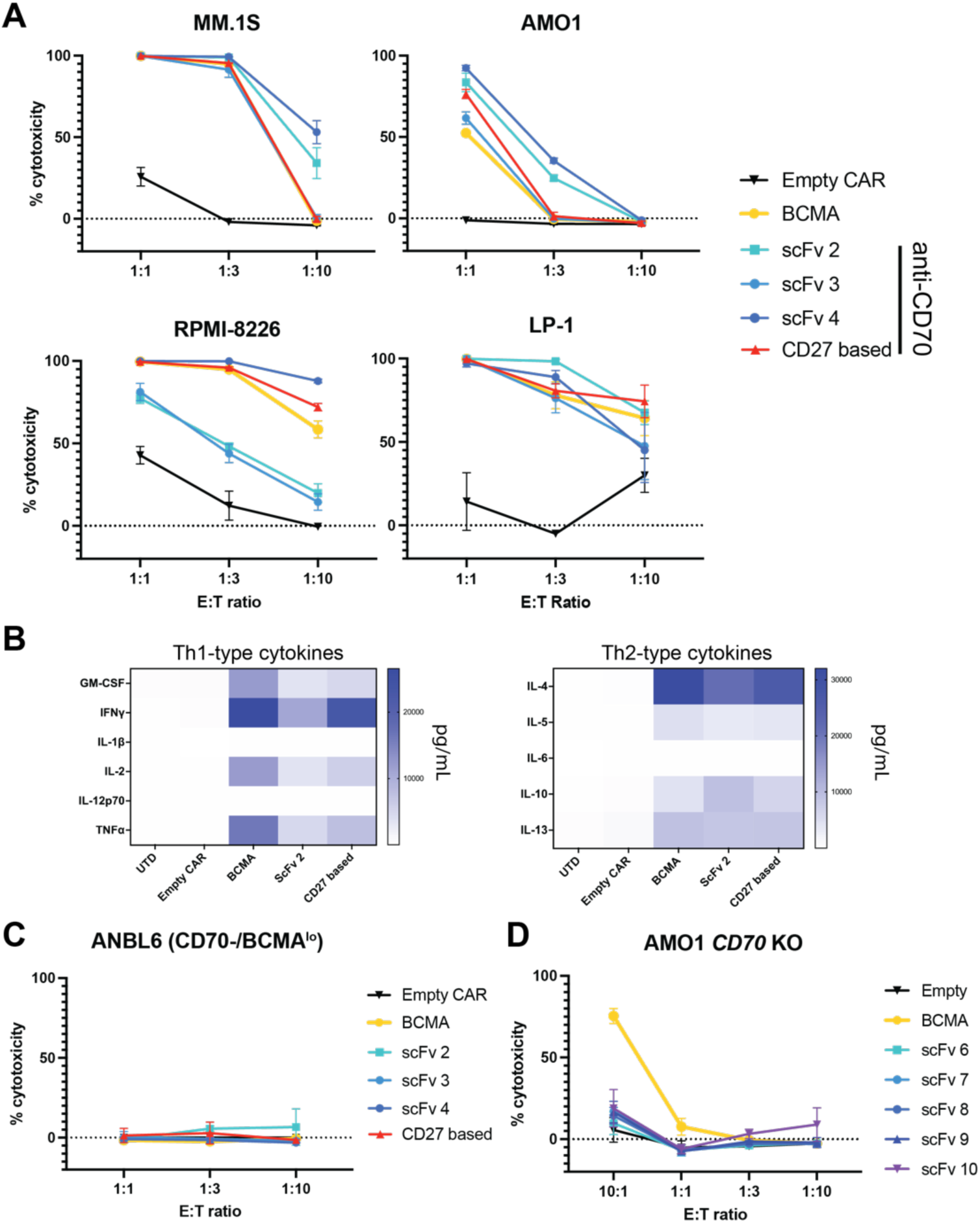
Both CD27-based and scFv-based CAR-Ts are effective versus myeloma models in vitro. **A.** In vitro cytotoxicity of FL CD27 ECD-based CARs and 3 scFv-based CARs versus several myeloma cell line models. **B.** Multiplexed cytokine analysis (Eve Biosciences) of CAR T supernatant, after overnight co-culture at 1:1 E:T with LP-1 cells. *n* = 3 technical replicates. **C-D.** *In vitro* cytotoxicity of noted CAR designs versus ANBL-6 myeloma cells (CD70 negative, low BCMA expression) (C) and AMO-1 cells with *CD70* knocked out by CRISPR/Cas9 (D). A, C, D: 18 hrs assay, lysis measured by luciferase at noted E:T ratios. *n* = 3 technical replicates, representative of 2 biological replicates from different T-cell donors. Mean +/− S.D. shown.

After overnight *in vitro* co-culture with LP-1 cells, flow cytometry for stem-ness markers indicated an increased percentage of effector memory T-cells in CD27-based CARs compared to other designs (**Fig. S4B**). Exhaustion markers revealed only modest differences between BCMA-targeting and CD27-based CAR-Ts (**Fig. S4C**). *In vitro* cytokine profiling after MM.1S co-culture showed increased release of several cytokines including TNF-a, IFN-g, and IL-2 for CD27-based vs. scFv 2-based anti-CD70 CAR-T, though BCMA CAR exceeded both anti-CD70 designs (**Fig. 3B**, **Fig. S4D**). Specificity testing versus the ANBL-6 MM line, which we found to lack CD70 and express very low BCMA (**Fig. S4E**), showed no activity of any of the tested CAR-Ts (**Fig. 3C**). CRISPR-based KO of *CD70* in an AMO-1 cell line also abrogated activity of all the anti-CD70 CARs, as expected, but not anti-BCMA CAR (**Fig. 3D**). Consistent with prior studies (*23*), anti-CD70 CAR-T constructs did not impact the function of hematopoietic progenitors in colony-forming assays (**Fig. S5A**). Taken together, these results initially demonstrate the activity of anti-CD70 CAR-T cells in MM preclinical models.

### CD27-based CAR-T cells display increased expansion in vivo compared to scFv CAR-T cells

These findings motivated us to perform an initial *in vivo* study of the three top-performing scFv-based CARs compared to CD27-based CAR. Anti-BCMA CAR-T was used as a positive control and “empty” CAR (same CAR backbone as anti-CD70 CARs but with no antigen recognition domain) was used as negative control. We intravenously implanted 1e6 luciferase-expressing MM.1S cells in NSG mice and then administered 5e6 CAR-T cells 7 days later. We were encouraged to find that all of the anti-CD70 CARs, as well as the anti-BCMA CAR, readily controlled the tumor burden in this model, supporting potential efficacy of anti-CD70 CAR-Ts in MM (**Fig. 4A-4B**).

**Figure 4.**
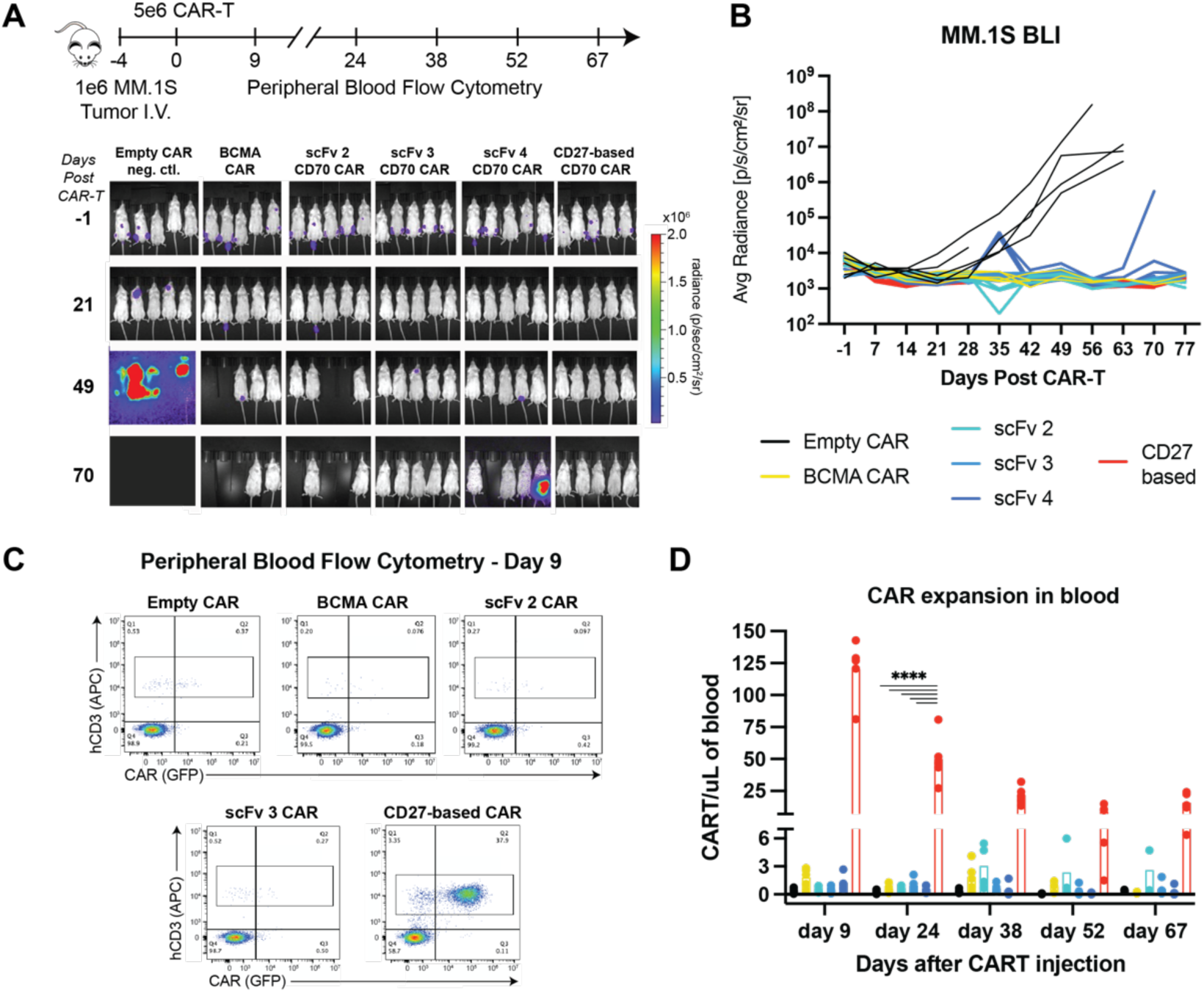
CD27-based CAR-Ts lead to dramatically increased expansion in vivo in an MM model. **A.** Murine study design and bioluminescent imaging. Disseminated MM.1S myeloma model implanted intravenously in NSG mice, *n* = 5/arm. Three anti-CD70 CAR designs are evaluated (scFv-2, -3, -4) compared to FL ECD CD27 CAR. Note that at later time points several mice in BCMA CAR, scFv 2 CAR, and scFv 4 CAR arms were required to be sacrificed without any evidence of tumor burden, likely due to development of graft-vs-host-disease (GvHD) based on murine symptoms. **B.** Quantified bioluminescence data illustrating tumor control for all anti-CD70 CARs. **C.** Representative flow cytometry plots of murine peripheral blood at Day 9. CAR Ts were quantified based on human CD3 (hCD3) and GFP positivity. **D.** Quantification of longitudinal CAR T expansion in murine peripheral blood. Split axis used to illustrate different scales of expansion for CD27 CAR versus others. Figure legend same as in B. p-value by 2-way ANOVA. ****p<0.0001. Mean +/− S.D. shown.

In parallel, we also monitored CAR-T expansion in murine peripheral blood by flow cytometry. Gating on human CD3+ T-cells at biweekly time points, we found that GFP+ scFv-based anti-CD70 CARs and anti-BCMA CAR-T comprised ~0.1-1% of all T cells (**Fig. 4C**). Remarkably, though, we found that the CD27-based CAR-T had ~80-100-fold increased expansion across all time points (**Fig. 4C-D**). Notably, a prior study of anti-CD70 CAR in AML models, using the full CD27 sequence fused to CD3z, also found a similar increased expansion phenotype compared to scFv-based CARs (*23*). However, in the prior AML study, this dynamic was thought to be due to different signaling incurred by the CD27 transmembrane and intracellular components. Here, we show that the CD27 ECD alone, fused to the identical CAR backbone as comparator scFv’s, can drive this dramatically increased expansion and persistence. Notably, for most clinically-tested CAR-Ts, such phenotypes appear highly favorable in terms of driving improved patient responses (*38, 39*). Together, these results therefore underscore the potential of optimized CD27-based CAR-Ts as a promising therapeutic strategy for MM patients with CD70+ tumors.

### Endogenous CD27 knockout in CAR-T cells does not eliminate robust expansion

In parallel, we also sought to investigate whether these CAR-T expansion differences could replicate in a different MM model *in vivo*. In this case we used the LP-1 model which carries the t(4;14) translocation, to recapitulate the MM subtype which may most frequently benefit from anti-CD70 therapy (**Fig. 1B**). In the LP-1 model, we again observed high efficacy and markedly increased *in vivo* expansion of CD27-based CAR-Ts compared to scFv-based CARs (**Fig. 5A-C**), replicating our findings in the prior MM.1S model.

**Figure 5.**
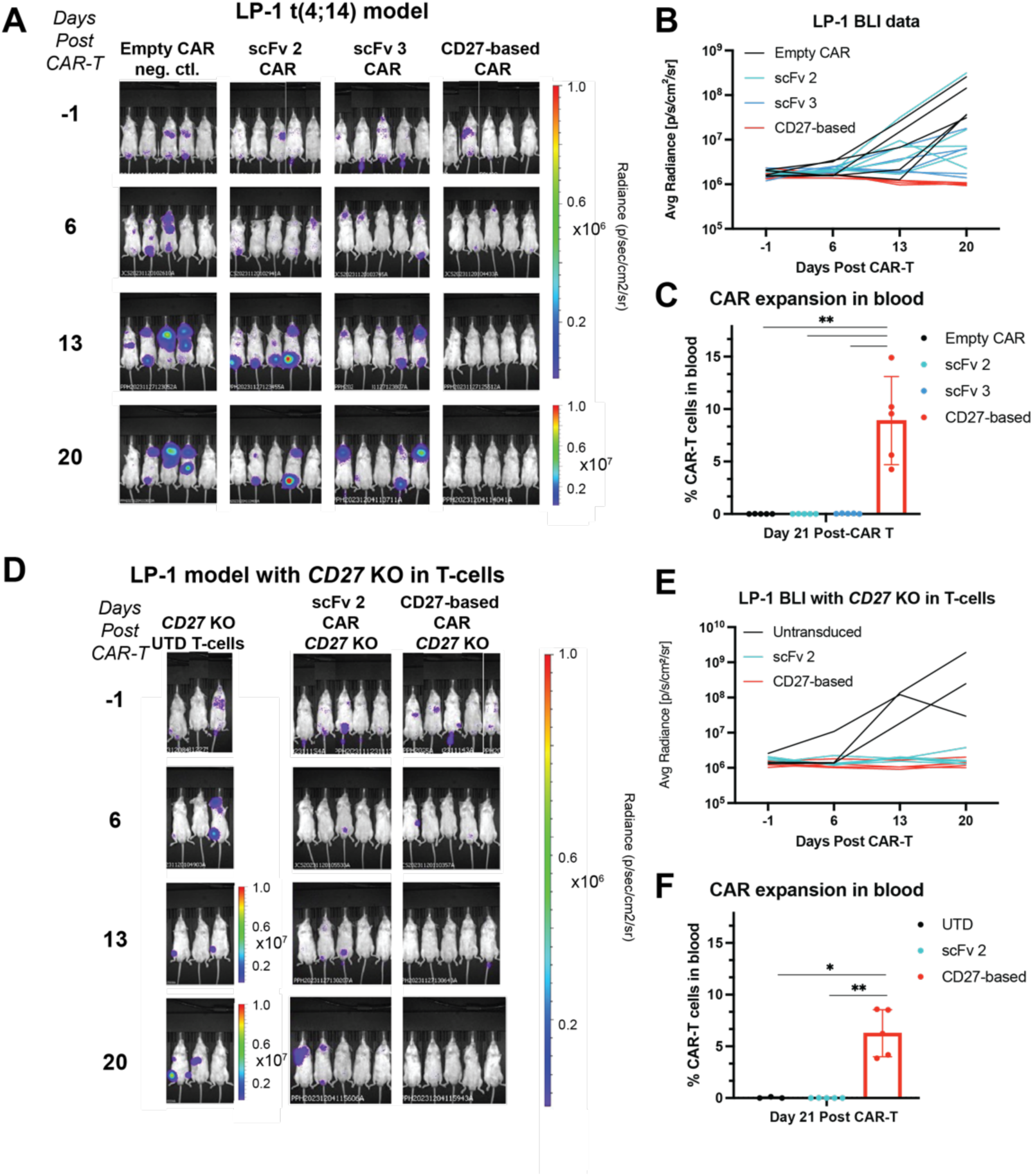
*CD27* knockout from T-cells does not abrogate expansion phenotype. **A.** The LP-1 myeloma model harboring t(4;14) translocation (1e6 tumor cells/mouse) was implanted intravenously in NSG mice and 5e6 CAR T cells implanted 5 days later. *n* = 5 mice/arm. Note increased scale for bioluminescence measurement at final time point. T-cells from a separate donor as used in Fig. 4. Study ended after Day 21 due to development of GvHD symptoms in several mice across arms. **B.** Bioluminescence from the study illustrating superior tumor control by CD27-based CAR compared to others. **C.** CD27-based CAR exhibits superior CAR-T expansion over scFv-based anti-CD70 CARs. Gating strategy same as in Fig. 5C. *p*-value by 2-way ANOVA. \*\**p*<0.01. **D.** *CD27* was knocked out donor T-cells (see Fig. S5 for flow cytometry validation) prior to transduction of each CAR, and then used in same intravenously-implanted LP-1 model as (C). Note increased radiance scale for UTD (untransduced) T-cell negative control at later time points. *n* = 5 mice/arm for scFv 2 and CD27-based CAR arms, *n* = 3 mice/arm for UTD. **E.** Quantification of bioluminescence shows continued tumor control by CD27-based CAR in this model. **F.** CD27-based CAR expansion in blood persists in this model despite CD27 KO from T cells. *p*-value by 2-way ANOVA. *p < 0.05, **p < 0.01. Difference in expansion between CARs from CD27 WT vs. *CD27* KO cells was non-significant (Fig. S5). Mean +/− S.D. shown.

We initially investigated why CD27-based CARs exhibit significantly increased expansion *in vivo*. One mechanistic hypothesis we explored is related to possible *cis* interactions of the CD27 ECD-based CARs with endogenous CD27 present on the surface of the CAR-Ts (**Fig. 5D**). In the LP-1 study, we therefore included two additional arms of the study, where in addition to *CD70* knockout to prevent fratricide, we also used Cas9 RNP to simultaneously knock out endogenous *CD27* from the CAR-T cells (**Fig. S5B**). We compared expansion of these “double KO” cells in the context of both scFv-based and CD27-based CARs. We again found that scFv-based CARs were only sparsely detectable in murine blood, similar to the single KO (*CD70* KO only) CARs, whereas CD27-based CARs were strongly expanded (**Fig. 5F**). Indeed, for CD27-based CARs, we found that *CD27* KO did not significantly abrogate the T-cell expansion phenotype (**Fig. S5C**). These findings therefore indicate that endogenous CD27 on CD27 ECD-based CAR-T cells is unlikely to be the critical factor relating to marked *in vivo* expansion.

### Identifying key transcriptional regulators of CD70 in myeloma

We next turned to the question of why *CD70* is specifically upregulated in high risk MM subtypes. Identifying mechanisms of upregulation could assist in identifying additional biomarkers predictive of response to CD70-based therapy, as well as predicting possible mechanisms of resistance to therapy due to antigen downregulation.

For the t(4;14) genomic subtype of MM, the known major oncogenic driver is overexpression of the methyltransferase *NSD2* (also known as *MMSET* or *WHSC1*) leading to a dysregulated transcriptional program (*40*). We therefore first asked whether NSD2 may play a role in the significant upregulation of *CD70* in t(4;14). Indeed, knockout of *NSD2* in two different t(4;14) models both led to decreased surface CD70 (**Fig. 6A**, **Fig. S6A**), supporting a functional role for this key epigenetic modifier in high CD70 expression in t(4;14) MM.

**Figure 6.**
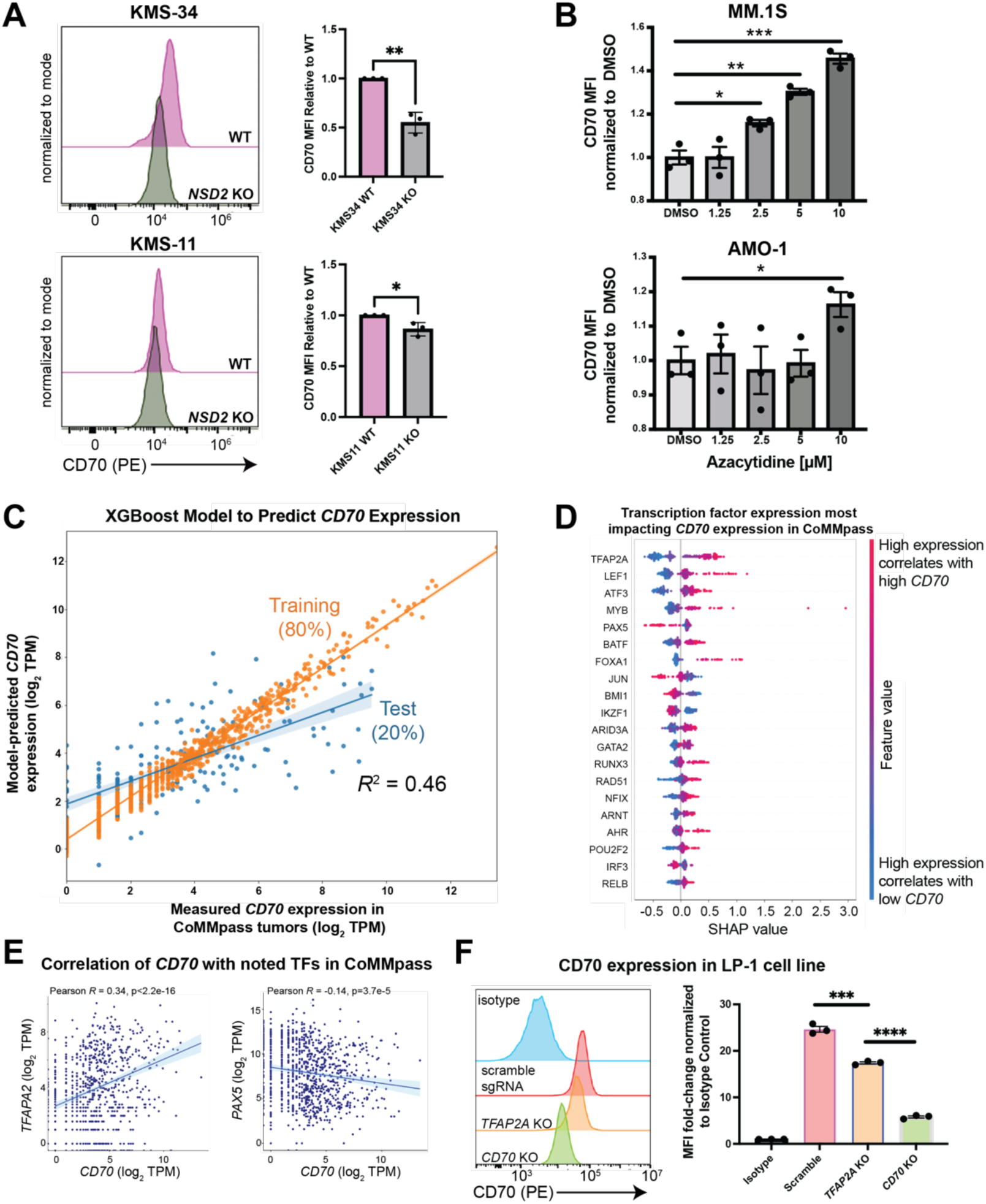
Identifying regulators of *CD70* expression in multiple myeloma. **A.** Knockout of the driver oncogene *NSD2* in t(4;14) myeloma cell lines KMS-11 and KMS-34 led to decreased surface CD70 expression by flow cytometry. *n* = 3 biological replicates. *p*-value by *t*-test. *p<0.05, **p<0.01. **B.** MM.1S or AMO.1 myeloma cells were treated with the indicated dose of azacytidine or 0.1% DMSO for 4 days, and surface CD70 assessed by flow cytometry 3 days later. *p*-value by 2-way ANOVA. \**p*<0.05, \*\**p*<0.01. *n* = 3 technical replicates. **C.** An eXtreme Gradient Boosting (XGB) model (see Methods) extracts features of transcription factor gene expression from CoMMpass (*n* = 776) that best-model *CD70* expression in patient tumors. 80% of data was used as a test set with 20% left out as a training set, with subsequent 5-fold cross validation (see Methods). *n* = 776 total samples included in the analysis. *R*^2^ value corresponds to Pearson correlation between model-predicted *CD70* expression in the test set and measured tumor *CD70* expression in CoMMpass. **D.** Shapley Additive Explanations (SHAP) analysis suggest expression of transcription factors most strongly impacting *CD70* expression levels in CoMMpass tumors. **E.** Correlation of *CD70* with *TFAP2A* and *PAX5* in the CoMMpass dataset. **F.** Surface CD70 by flow cytometry on LP-1 cells measured after Cas9 RNP nucleofection with sgRNA targeting *TFAP2A*, *CD70*, or scramble control. Flow cytometry mean fluorescence intensity shown as fold-change versus isotype control. *p*-value by *t*-test. \*\*\**p*<0.005, ****p<0.001. *n* = 3 technical replicates, representative of 2 biological replicates. Mean +/− S.D. shown.

Other groups previously established in AML models that treatment with the DNA methyltransferase inhibitor azacytidine could lead to increased CD70 expression (*16, 22*). We also recapitulated this finding in MM models (**Fig. 6B**), suggesting a possible future co-treatment strategy with CD70-targeted immunotherapies for this malignancy.

These findings encouraged us to search for other epigenetic and transcriptional regulators of *CD70* in MM. We used a strategy previously described by our group (*41*), analyzing both primary MM sample ATAC-seq data (*42*) and ENCODE ChIP-seq data, to together predict transcription factors (TFs) that may bind to the *CD70* locus (**Fig. S6B**). We then took this list of 84 transcription factors and correlated TF expression with *CD70* expression across all 776 patient tumor samples in CoMMpass. Using these correlations, we then built a machine learning classifier via eXtreme Gradient Boosting (XGBoost) (*41*). This model enabled us to determine the relative contribution of expression of each TF with respect to *CD70* expression. Using 80% of the CoMMpass data to train the classifier and the remainder as a test set, we found that the TF-based classifier was moderately predictive (*R*^2^ = 0.46) of *CD70* expression in MM tumors (**Fig. 6C**).

These results suggest that *CD70* expression can be at least partially predicted based on expression of associated transcriptional regulators. SHAP analysis further delineated TFs with the greatest weight in the XGBoost model including potential positive regulators *TFAP2A* (AP-2), *LEF1*, and *ATF3*, and top negative regulator *PAX5* (**Fig. 6D,E**). To functionally validate these findings, we used a Cas9 RNP approach to knock out *TFAP2A* in the LP-1 t(4;14) model. The positive control of *CD70* knockout, as expected, led to significant reduction of surface CD70 when compared to non-targeting control (**Fig. 6F**, **Fig. S7A**). *TFAP2A* KO also led to decreased surface CD70, albeit by a smaller magnitude than *CD70* KO (**Fig. 6F**). This result is expected per our model: while these TFs are predicted to contribute to *CD70* transcript levels, they each only play a partial, not complete, role in its expression.

As the top predicted negative regulator, *PAX5* could not be readily probed as it does not have detectable expression in any of the commonly used MM cell line models (**Fig. S7B**). However, *PAX5*, a TF closely linked to B-cell identity (*43*), is most highly expressed in the t(11;14) subtype of MM (**Fig. S7C**). Intriguingly, we found this standard-risk subtype to have the lowest *CD70* expression (**Fig. 1B**) and we have also recently characterized t(11;14) to broadly exhibit a B-cell like phenotype (*44*). These results raise the hypothesis that a more B-cell-like profile may suppress CD70 expression.

STRING analysis indicates that nearly all transcription factors that our model predicts to play a role in *CD70* regulation form a tight network of interactions with potential inter-regulation (**Fig. S7D**). Taken together, these findings identify possible relevant transcriptional regulators of *CD70* in high risk MM and point to specific patterns of gene expression and activation that drive expression of this immunotherapy target.

### Dual-targeting, bicistronic CARs vs. CD70 and BCMA to avoid antigen escape

Our work above suggests that CD70 may be a promising cell surface target for high-risk tumors relapsing after BCMA-targeting immunotherapies. Another potential strategy could be a first-line cell therapy that simultaneously targets both BCMA and CD70 to achieve even deeper and more durable remissions in high-risk patients. Dual-targeting CAR-Ts have been proposed to be effective in tumors with heterogeneous expression of the targeted surface antigens, or to avoid tumor resistance due to downregulation of one antigen (“antigen escape”) (*45*). Similar dual-targeting strategies have been described for BCMA and GPRC5D (*46*). To this end, we developed a proof-of-principle dual-targeting CAR-T using a bicistronic construct with both CARs separated by a T2A element (**Fig. 7A-B**, **Fig. S8A**). In initial *in vitro* studies, we showed that this CAR could successfully eliminate LP-1 cells with either *TNFSRF17A* (which encodes BCMA) or *CD70* knocked out (**Fig. S8B**), mimicking antigen escape conditions for these targets. Demonstrating specificity, this dual-targeting CAR-T was ineffective against LP-1 cells with both genes knocked out (**Fig. 7C**). Though further work is clearly required to generate a true therapeutic candidate, here we aim to described a proof-of-principle construct as a possible complementary strategy for future clinical translation of CD27-based CAR-Ts.

**Figure 7.**
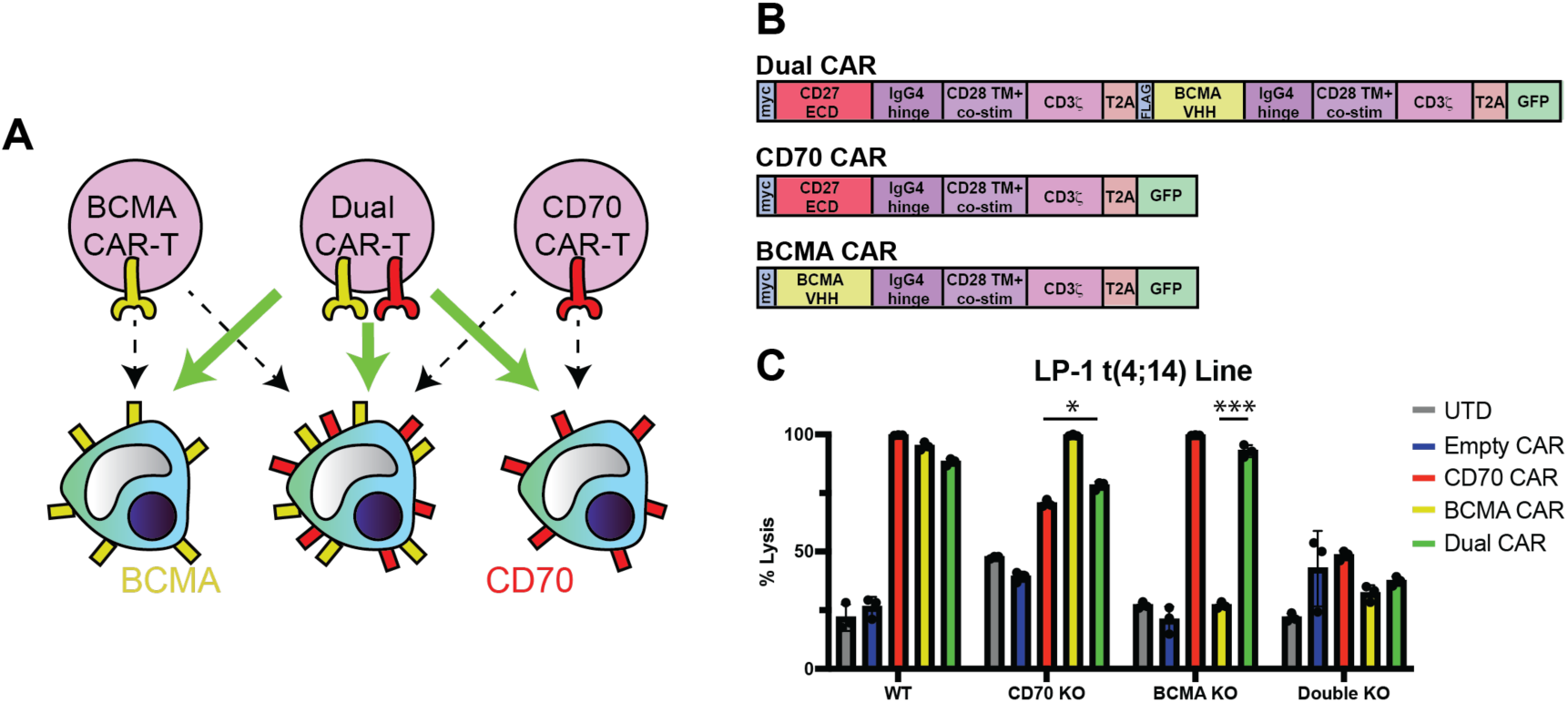
Proof-of-principle dual-targeting CAR-T against both CD70 and BCMA can potentially overcome antigen escape. **A.** Illustration of dual-targeting anti-CD70 + anti-BCMA CAR-T to eliminate myeloma cells that do not display CD70 or BCMA on the cell surface. **B.** Design of bi-cistronic dual-targeting and mono-cistronic single-targeting CAR constructs. **C.** Cytotoxicity of dual-targeting CAR-T versus LP-1 cell (WT, CD70 KO, or BCMA KO; see Fig. S8); *n* = 3 technical replicates, 24 hr assay, 3:1 E:T. *p*-value by two-tailed *t*-test. Comparisons shown for dual CAR vs. each single-targeting CAR under corresponding single gene knockout conditions. \**p*<0.05, \*\*\**p*<0.005. Mean +/− S.D. shown.

## Discussion

Here we demonstrate that CD70, a known immunotherapy target in several malignancies with a highly favorable “on target, off tumor” toxicity profile, is also a promising and viable cellular therapy target for several subtypes of high risk MM. We further use structure-guided design to optimize a “natural ligand”-based CAR-T vs. CD70 with greatly improved *in vivo* expansion and persistence compared to scFv-based CARs. Epigenetic analysis reveals potential transcriptional regulators of CD70 and further elucidates tumor signatures which may confer utility for CD70-targeting therapeutics.

This work describes a potential new therapeutic option for MM patients most in need in the BCMA CAR-T era. Though GPRC5D-targeting CAR-Ts have shown significant promise as a next-line treatment, this approach still does not appear curative (*9*). In addition, on target, off tumor toxicity to nail beds and skin are known, though clinically manageable, complications of this approach (*47*). As MM transitions to become a chronic disease (*48*), identifying additional next-line therapies, particularly for high-risk patients, will continue to be an important part of patient management. CD70-targeting CAR-Ts, with potential for strong efficacy but low toxicity, could thus have significant scope for clinical utility. In addition, given that CD70 is expressed minimally on normal immune cells (*15*), it is possible that CD70-targeting CAR-Ts may avoid long-term immune complications such as B-cell or plasma cell aplasia as seen with BCMA CAR-Ts (*49*).

From a CAR design perspective, our work offers important insight into the impact of the antigen recognition element in the anti-CD70 CAR. Prior work from Leick et al. (*22*) found that eliminating the native CD27 transmembrane sequence from the CAR could increase anti-AML potency by avoiding proteolysis of the CD27 ECD. In parallel, Sauer et al. (*23*) found that a full-length CD27 sequence (including CD27 extracellular, transmembrane, and intracellular domains) could lead to increased *in vivo* expansion in AML models over scFv-based CARs. However, in that latter study it was not clear if this increased expansion was caused by CD27-based signaling differences, or difference in the antigen recognition domain. Here, we definitively show that greatly increased *in vivo* expansion occurs as a function of the CD27 ECD alone, even with an identical CAR backbone architecture to scFv-based designs. We also showed that this increased expansion was not driven by interaction of the CAR-fused CD27 ECD with endogenous CD27 present on the T-cell surface. While further investigation will be required, this outcome suggests that the signal transduction through the CAR itself has significantly altered dynamics as a function of either CD27- or scFv-based engagement. This observation could have important implications for other natural ligand-based CARs, which in general are highly promising given a fully human sequence with potential for low immunogenicity when compared to antibody fragments (*32*).

In terms of limitations of our study, CD70 is not upregulated on all subtypes of high risk MM. While we find clear association of CD70 with gain of Chr1q, present in ~40% of MM patients, and t(4:14), present in ~15% of patients, another major subtype of high risk disease, patients with deletion of Chr17p or biallelic *TP53* loss (~10% of patients) (*27*), do not show increased *CD70* expression. In addition, even on CD70+ tumors, this antigen may be expressed at lower levels on malignant plasma cells than BCMA. Identifying even further-optimized CD27-based CAR designs may be important to fully eliminate tumor cells in patient tumors with low antigen density. Future work will evaluate whether natural ligand CARs targeting CD70 can be enhanced for low antigen recognition using protein computational design strategies. Alternatively, co-treatment strategies with azacytidine or other small molecules could be used to increase tumor antigen density and enhance potency of anti-CD70 CAR-Ts. Further characterization of critical transcriptional and regulatory networks, as we begin to delineate here, could reveal such additional strategies to drive CD70 upregulation in MM tumors. Though additional evaluation is required, dual-targeting CARs against both CD70 and BCMA could also address issues of surface protein heterogeneity while standing as a potent yet safe cell therapy.

Notably, our results support the notion that even current CD70-targeting CAR-Ts under clinical development for other indications (for example, NCT05948033, NCT02830724, NCT05947487, NCT04502446) could consider immediate inclusion of patients with CD70+ MM tumors. Even though most of these current anti-CD70 therapies employ scFv-based designs, many MM patients could still benefit, and clinical data from these studies could be leveraged to inform future studies employing more potent, optimized CD27-based CARs.

## Acknowledgements

We would like to acknowledge contributions of the staff of the UCSF clinical flow cytometry laboratory and patients and their families for participating in research.

## Author Contributions

C.K., A.I., B.P.E., A.P.W. designed and conceived the study. C.K. and A.P.W. wrote the manuscript with input from all authors. C.K., B.P.E., A.I., A.K., N.C., H.J., J.H., T.R., A.B., A.S., and D. D.-R. performed experiments and analyzed data. S.R., D.G.-A., G.W., and K.M.K.K. performed experiments. B.P.E., N.A., H.G., Y-H.T.L., E.R., A.J., W.J.K., and B.G.B. analyzed patient data and datasets. F.S., P.P., J.A.C.S., O.Z., I.T., and V.S. performed murine studies. M.S., L.H.B., J.D.L., and E.S. participated in experimental design and funding acquisition. R.D. and T.K. performed structural protein analysis.

## Accession numbers/Data Availability

AML and myeloma cell line surface proteomics datasets: Raw and processed datasets available at PRIDE repository under accession number **PXD048492**.

*Reviewer access details:*

Username: Password: DHshh9Q1

## Code Availability

R scripts used for CoMMpass analysis and machine learning model available at: https://github.com/BonellPatinoE/CD70-in-Multiple-Myeloma.git

## Materials and Methods

### Human Samples and Ethics

Patient samples were collected under University of California, San Francisco (UCSF) Institutional Review Board-approved protocols and in accordance with the Declaration of Helsinki. All murine studies were conducted under UCSF Institutional Animal Care and Use Committee-approved protocols.

### Cell lines

Human cell lines were authenticated by short tandem repeat (STR) analysis and routinely tested for mycoplasma contamination. Cells were grown in RPMI 1640 medium supplemented with 20% fetal bovine serum (FBS) and 100 U/mL penicillin-streptomycin. Cell lines used were RPMI-8226, MM.1S (originally obtained from ATCC); AMO-1, LP-1 (originally obtained from DSMZ); KMS-11, KMS-34 (originally obtained from JCRB).

### Molecular Cloning and DNA Plasmids

Genes encoding the different anti-CD70 (either scFv or CD27 and their truncations) sequences were synthesized as gene fragments from Twist Biosciences (South San Francisco, CA). DNA fragments were then cloned into a lentiviral expression vector with a Gibson assembly product (NEB, E2611S). DNA sequencing was performed to confirm accuracy of the vector. This final construct was then expressed in NEB 5-alpha Competent E. coli (High Efficiency) (NEB, C2987H) and Stbl3 Competent E. coli (Macro Lab, UC Berkeley, CA). DNA was isolated using either QIAGEN Plasmid Plus Midi Kit or 28 QiaPrep Spin Miniprep Kit (Qiagen, 27104).

### Lentiviral Vector Production

Lenti-X 293T cells (Takara, Cat # 632180) were transfected with each CAR expression plasmid and Mirus Bio™ TransIT™-Lenti Transfection Reagent (Cat # MIR6600). For Bicistronic dual-targeting CAR-T, Invitrogen™ Lipofectamine™ 3000 Transfection Reagent (Cat # L3000015) was used along with ViralBoost Reagent (Alstem Cell Advancements, Richmond, CA. Cat # VB100). Lenti-X 293T cells were cultured for 2-3 days, and then lentivirus was harvested and concentrated using Lenti-X Concentrator (Takara Bio, 631232).

### CAR T-cell Production and Expansion

Primary human T cells were purified from either 1) the leukoreduction filter products of anonymous healthy blood donors from Vitalant (San Francisco, CA) under an institutional review board–exempt protocol in accordance with the U.S. Common Rule (Category 4) or 2) de-identified donor leukapheresis products obtained from StemCell Technologies. CD3+ T-cell populations were isolated using RosetteSep Human T Cell Enrichment Cocktails (Stemcell Technologies, 15023 and 15022) and EasySep Human CD4 T Cell Iso Kit (Stemcell Technologies, 17952) and EasySep Human CD8 T Cell Iso Kit (Stemcell Technologies, 17953). CD3+ T-cells were thawed, cultured separately in media overnight, and then cells were counted the following day. T cells were cultured in OpTmizer medium with CTS supplement (Thermo Scientific, A1048501) supplemented with 5% human AB serum (HP1022; Valley Medical), GlutaMAX (Gibco™. Cat # 35050061), and 100 U/mL penicillin/streptomycin (Fisher Scientific, 15-140-122) and were passaged every 2 days. For expansion, T cells were stimulated with CD3/CD28 Dynabeads (11131-D; Thermo Fisher Scientific) according to the manufacturer’s instructions (20 μL of beads per 1 million T-cells) for 5 days and grown in the presence of recombinant interleukin-7 (PeproTech, 200-07) and interleukin-15 (PeproTech, 200-15) at 10 ng/mL. Transduction with CAR lentivirus was performed 1 day after the start of bead stimulation. After the removal of CD3/CD28 activation beads, transduction efficiency was assessed by flow cytometry. CAR-T cells were labeled with intracellular GFP, so the percentage of GFP+ cells determined the percentage of CAR-T cells generated.

### Flow Cytometry

Cells were resuspended in FACS buffer (2% FBS in D-PBS) and stained with antibodies for 30-60 min at 4C, washed with FACS buffer, and then resuspended in FACS buffer. Samples were analyzed using either a Cytoflex Flow Cytometer (Beckman Coulter, Beckman Coulter Navios Flow Cytometer) or FACSAria-Fusion or FACSAria III flow cytometer (BD Biosciences). Data analysis was done using FlowJo software, v10.10.0. Compensation was performed with UltraComp eBeadsTM Compensation Beads. Invitrogen. Ref: 01-2222-42. Lot #: 2775954. For primary sample analysis, gating strategy used was FSC-A/SSC-A for lymphocyte population, single cells gated in SSC-A/SSC-H and then myeloma cell population was gated, either gating cells CD19−/CD138+ or CD45-/CD19−/CD138+/CD38+ to determine CD70 and BCMA expression.

### Antibodies

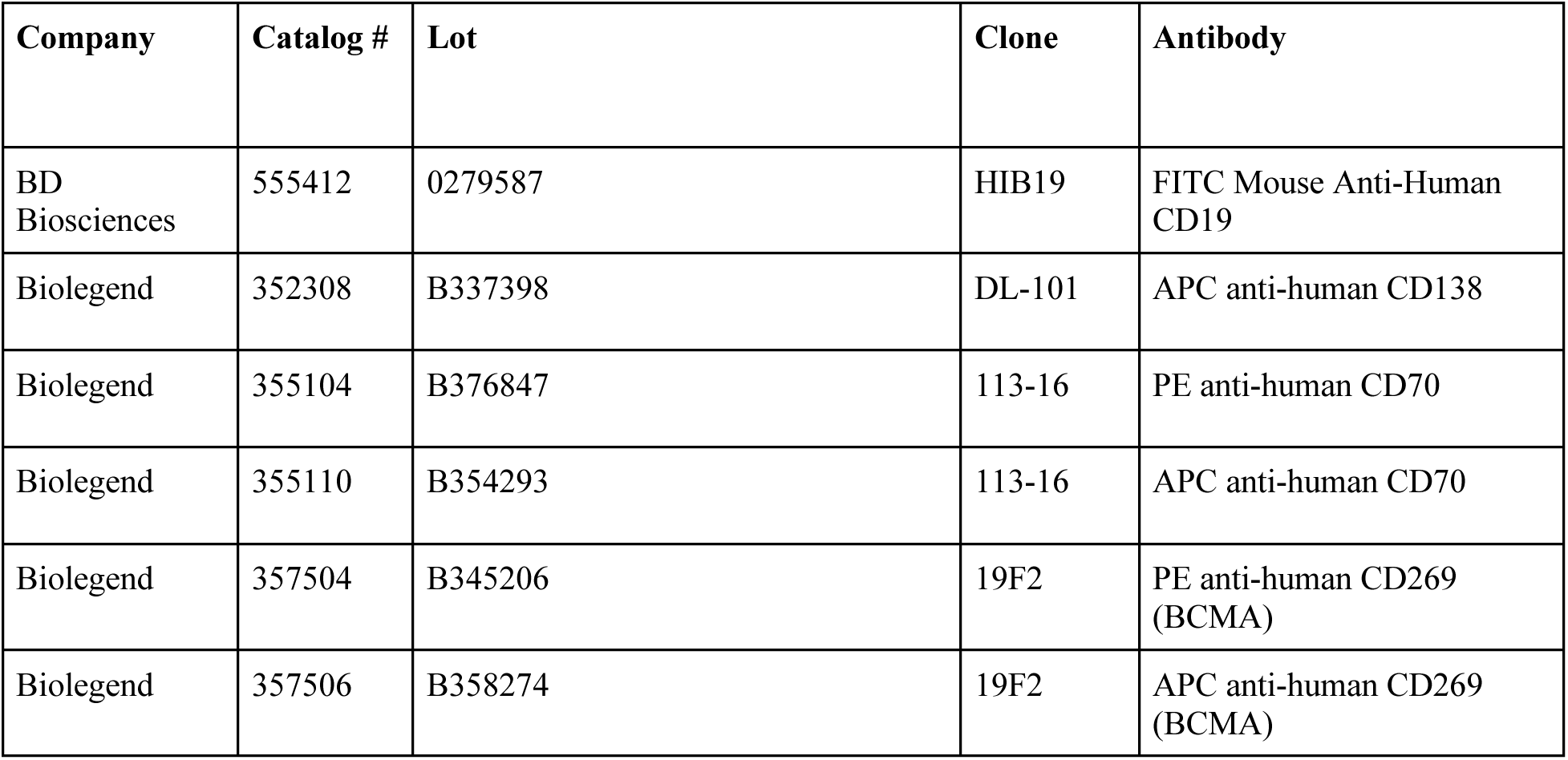
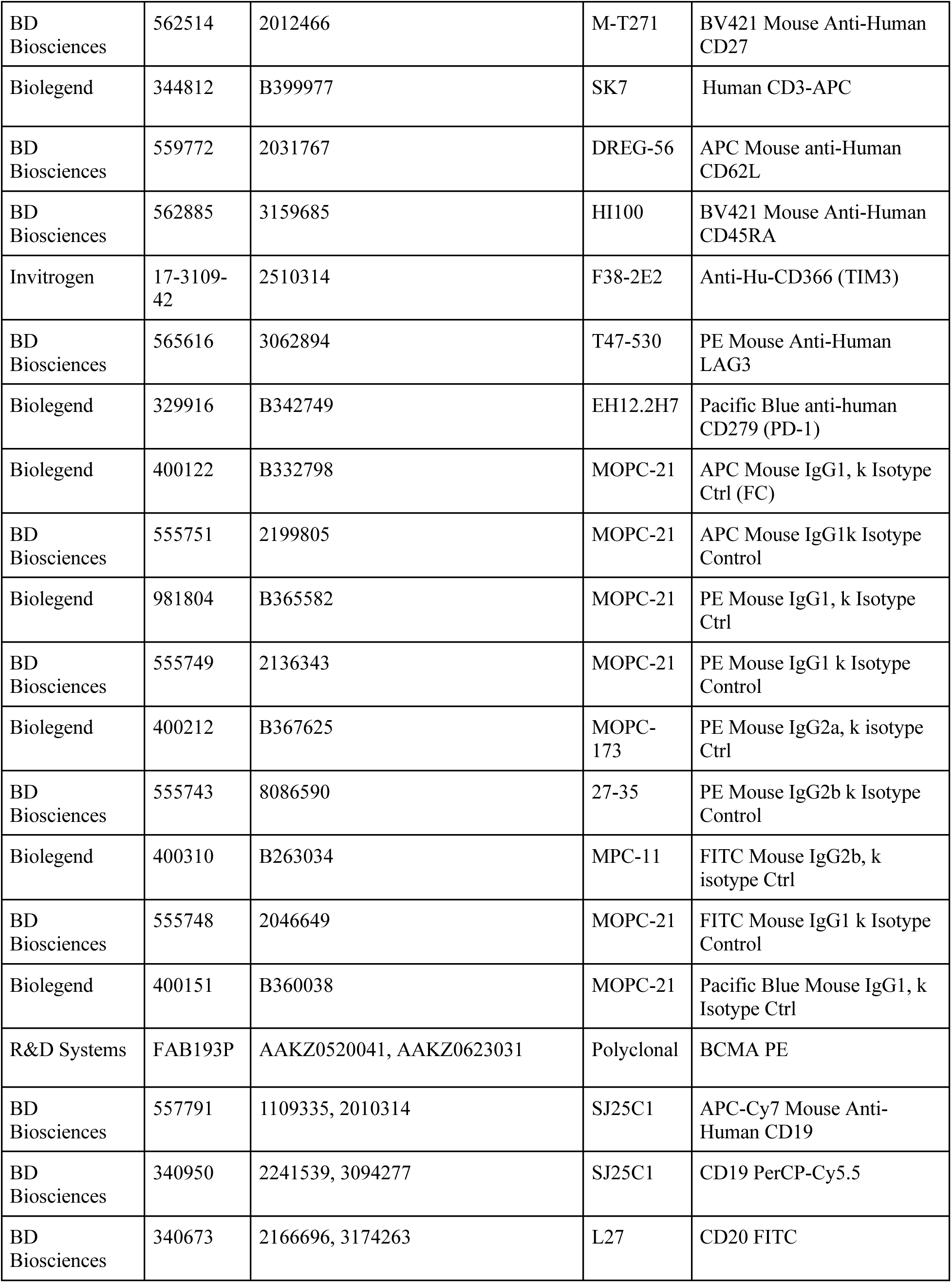
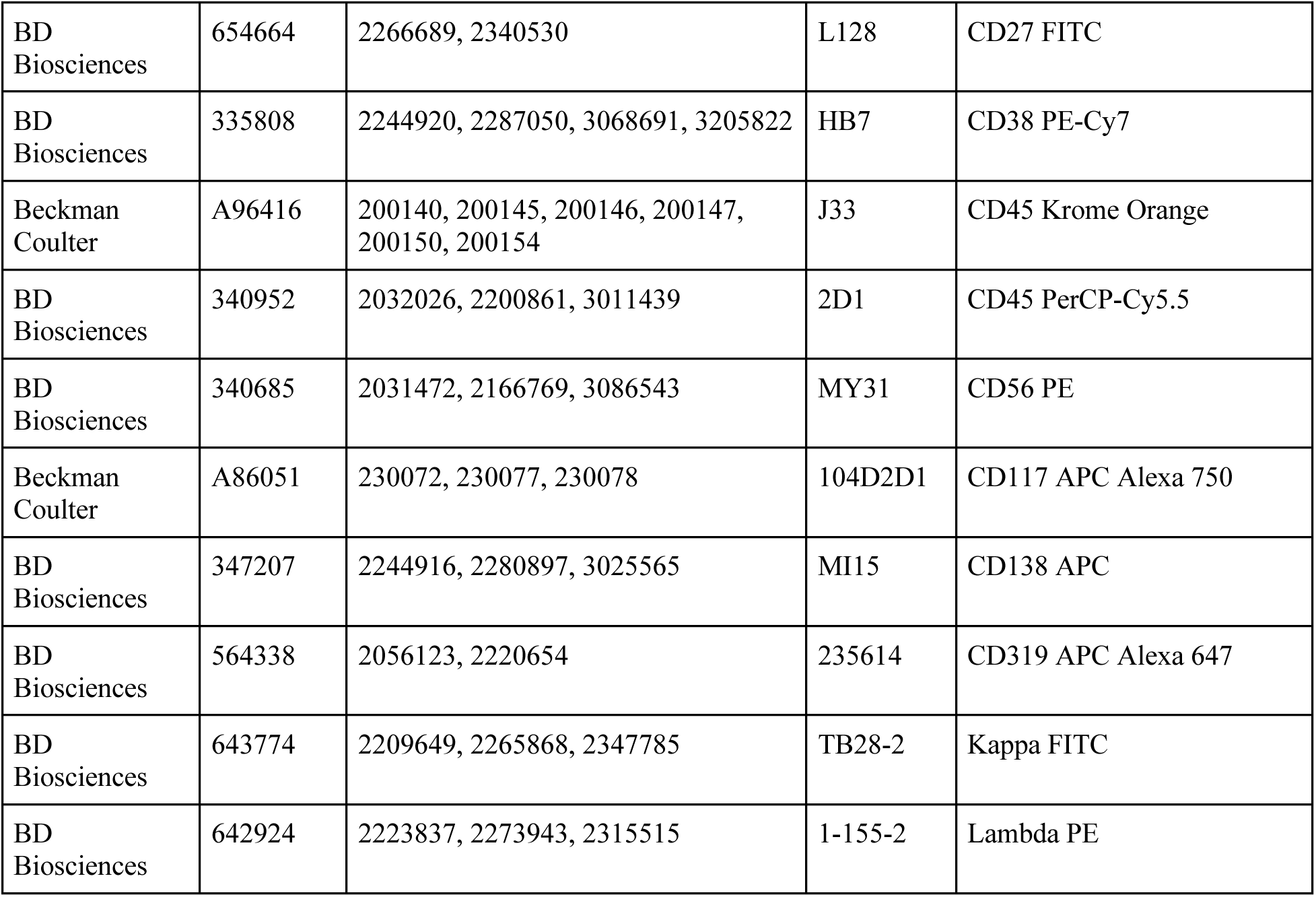

| Company | Catalog # | Lot | Clone | Antibody |
| --- | --- | --- | --- | --- |
| BD Biosciences | 555412 | 0279587 | HIB19 | FITC Mouse Anti-Human CD19 |
| Biolegend | 352308 | B337398 | DL-101 | APC anti-human CD138 |
| Biolegend | 355104 | B376847 | 113-16 | PE anti-human CD70 |
| Biolegend | 355110 | B354293 | 113-16 | APC anti-human CD70 |
| Biolegend | 357504 | B345206 | 19F2 | PE anti-human CD269 (BCMA) |
| Biolegend | 357506 | B358274 | 19F2 | APC anti-human CD269 (BCMA) |
| BD Biosciences | 562514 | 2012466 | M-T271 | BV421 Mouse Anti-Human CD27 |
| Biolegend | 344812 | B399977 | SK7 | Human CD3-APC |
| BD Biosciences | 559772 | 2031767 | DREG-56 | APC Mouse anti-Human CD62L |
| BD Biosciences | 562885 | 3159685 | HI100 | BV421 Mouse Anti-Human CD45RA |
| Invitrogen | 17-3109-42 | 2510314 | F38-2E2 | Anti-Hu-CD366 (TIM3) |
| BD Biosciences | 565616 | 3062894 | T47-530 | PE Mouse Anti-Human LAG3 |
| Biolegend | 329916 | B342749 | EH12.2H7 | Pacific Blue anti-human CD279 (PD-1) |
| Biolegend | 400122 | B332798 | MOPC-21 | APC Mouse IgG1, k Isotype Ctrl (FC) |
| BD Biosciences | 555751 | 2199805 | MOPC-21 | APC Mouse IgG1k Isotype Control |
| Biolegend | 981804 | B365582 | MOPC-21 | PE Mouse IgG1, k Isotype Ctrl |
| BD Biosciences | 555749 | 2136343 | MOPC-21 | PE Mouse IgG1 k Isotype Control |
| Biolegend | 400212 | B367625 | MOPC-173 | PE Mouse IgG2a, k isotype Ctrl |
| BD Biosciences | 555743 | 8086590 | 27-35 | PE Mouse IgG2b k Isotype Control |
| Biolegend | 400310 | B263034 | MPC-11 | FITC Mouse IgG2b, k isotype Ctrl |
| BD Biosciences | 555748 | 2046649 | MOPC-21 | FITC Mouse IgG1 k Isotype Control |
| Biolegend | 400151 | B360038 | MOPC-21 | Pacific Blue Mouse IgG1, k Isotype Ctrl |
| R&D Systems | FAB193P | AAKZ0520041, AAKZ0623031 | Polyclonal | BCMA PE |
| BD Biosciences | 557791 | 1109335, 2010314 | SJ25C1 | APC-Cy7 Mouse Anti-Human CD19 |
| BD Biosciences | 340950 | 2241539, 3094277 | SJ25C1 | CD19 PerCP-Cy5.5 |
| BD Biosciences | 340673 | 2166696, 3174263 | L27 | CD20 FITC |
| BD Biosciences | 654664 | 2266689, 2340530 | L128 | CD27 FITC |
| BD Biosciences | 335808 | 2244920, 2287050, 3068691, 3205822 | HB7 | CD38 PE-Cy7 |
| Beckman Coulter | A96416 | 200140, 200145, 200146, 200147, 200150, 200154 | J33 | CD45 Krome Orange |
| BD Biosciences | 340952 | 2032026, 2200861, 3011439 | 2D1 | CD45 PerCP-Cy5.5 |
| BD Biosciences | 340685 | 2031472, 2166769, 3086543 | MY31 | CD56 PE |
| Beckman Coulter | A86051 | 230072, 230077, 230078 | 104D2D1 | CD117 APC Alexa 750 |
| BD Biosciences | 347207 | 2244916, 2280897, 3025565 | MI15 | CD138 APC |
| BD Biosciences | 564338 | 2056123, 2220654 | 235614 | CD319 APC Alexa 647 |
| BD Biosciences | 643774 | 2209649, 2265868, 2347785 | TB28-2 | Kappa FITC |
| BD Biosciences | 642924 | 2223837, 2273943, 2315515 | 1-155-2 | Lambda PE |

### CoMMpass relapsed vs. new diagnosis analysis

Differential gene expression analysis was performed on RNAseq data from primary MM samples in the MMRF CoMMpass study (release IA14). The “HtSeq_Gene_Counts.txt” file was loaded and processed through DESeq2 (v1.34.0) in R (v4.1.2). First, we filtered for (50) patients with RNAseq data at both the initial diagnosis and first relapse. Second, we performed a paired analysis for differential gene expression between the two visits. Finally, we filtered for cell surface proteins using annotations from the literature (*50*) and visualized the results in a volcano plot.

### CoMMpass cytogenetic subtype transcriptional analysis

Differential gene expression analysis based on cytogenetic risk was performed on RNAseq data from primary MM samples from the Multiple Myeloma Research Foundation (MMRF) CoMMpass study (release IA19), which are available to registered users through the MMRF Researcher Gateway (registration information available at https://mmrf.formstack.com/forms/research_gateway_registration). The RNAseq data files were loaded and processed through DESeq2 (v1.34.0) (*51*) in R (v4.1.2), ‘apeglm’ for LFC shrinkage (*52*) and “gprofiler” (v0.2.2) for gene list functional enrichment analysis and namespace conversion (*53*). Using clinical information in CoMMpass, we performed a paired analysis for differential gene expression between ISS risk and *CD70* expression, matching patient ID and ISS stage determined in the clinical features data file. Cytogenetic risk features were determined from associated cytogenetics data set files to classify samples as positive or negative for specific rearrangements based on *CD70* expression. A similar approach was used for *PAX5* analysis. For survival analysis, Kaplan-Meier curves were made using survminer (0.4.9) and survival (v3.5-7) packages in R (v4.1.2) comparing top 20% and bottom 20% of *CD70* expression among patients of MMRF CoMMpass data set and defining survival events and time points. The correlation plots comparing *CD70* expression with predicted transcription factors determined as transcriptional regulators of *CD70* were made using ggscatter function in ggpubr package (version 0.6.0). The code used for our analysis is available at our repository here (https://github.com/BonellPatinoE/CD70-in-Multiple-Myeloma.git).

### CAR-T Cell Production and Expansion

Primary human T cells were purified from either 1) the leukoreduction filter products of anonymous healthy blood donors from Vitalant (San Francisco, CA) under an institutional review board–exempt protocol in accordance with the U.S. Common Rule (Category 4) or 2) de-identified donor leukapheresis products obtained from StemCell Technologies. CD3+ T-cell populations were isolated using EasySep Human T Cell Isolation Kit (StemCell Technologies, Cat# 17951, lOT #1000153615). CD3+ T-cells were thawed, cultured separately in media overnight, and then cells were counted the following day. T cells were cultured in OpTmizer medium with CTS supplement (Thermo Scientific, A1048501) supplemented with 5% human AB serum (HP1022; Valley Medical), GlutaMAX (Gibco™. Cat # 35050061), and 100 U/mL penicillin/streptomycin (Fisher Scientific, 15-140-122) and were passaged every 2 days. For expansion, T cells were stimulated with CD3/CD28 Dynabeads (11131-D; Thermo Fisher Scientific) according to the manufacturer’s instructions (20 μL of beads per 1 million T-cells) for 5 days and grown in the presence of recombinant interleukin-7 (PeproTech, 200-07) and interleukin-15 (PeproTech, 200-15) at 10 ng/mL. Transduction with CAR lentivirus was performed 1 day after the start of bead stimulation. After the removal of CD3/CD28 activation beads, transduction efficiency was assessed by flow cytometry. CAR-T cells were labeled with intracellular GFP, then the percentage of GFP+ cells determined the percentage of CAR-T cells generated.

### CD27 Truncation Design Generation and Production

Six truncated CD27 natural ligand CARs were generated to probe the importance of the first cysteine-rich domain (CRD-1) of CD27 for the function of a CD27 natural ligand-based CAR against CD70-positive targets. These 6 truncated designs, denoted from T1 to T6, represent increasing lengths of amino acids removed from the N-terminal end of CRD-1, with the shortest overall binder, T6, having CRD-1 removed entirely. The amino acid sequence of the binding region for each truncated CAR is listed as follows, using the amino acid numbering scheme from PDB:7KX0 (Liu et al., 2021) (*34*): T1: Gly 46-Pro 124; T2: Gly 46-Asp 121; T3: Phe 48-Pro 124; T4: Phe 48-Asp 121; T5: Leu 49-Pro 124; T6: Pro 64-Pro 124.

Constructs were designed using Snapgene (version 7.1.1) and Benchling. DNA fragments containing the truncated CAR binder sequences with 40-base pair target vector overlap regions were ordered from Twist Bioscience, then were incorporated into a 2nd generation CAR lentiviral backbone under the SFFV promoter using NEBuilder HiFi DNA Assembly kit following the manufacturer’s standard protocol. The overall CAR framework included a CD8α signal peptide, followed by the truncated CD27 binder, mutated IgG4 hinge domain(*54*), CD28 transmembrane domain, CD28 costimulatory domain, and CD3ζ signaling domain.

### In vitro cytotoxicity assays

MM.1S, LP-1, AMO-1, and RPMI-8226 MM cell lines were engineered to stably express luciferase using lentiviral transduction. For each cell line, 50,000 MM cells were seeded in 50 uL of T-cell media into a white 96 well plate (Greiner Bio-One). Subsequently, CAR-T cells were added at four different effector:target ratios (3:1, 1:1, 1:3, and 1:10) in 50uL of T-cell media with *n*=3 replicates per ratio and CAR-T construct. This coculture was incubated at 37C for 18-24 hours. The next day, 100uL of D-Luciferin (Gold Biotechnology, LUCK-1G) was added to each well to a final concentration of 375 μg/ml, followed by luminescence detection using GloMax Explorer (Promega). The bioluminescence readings were averaged amongst the *n*=3 technical replicates for each construct and ratio. Bioluminescence readings were normalized to the maximum and minimum values for each cell line to create a cytotoxicity scale of 0-100. Assays were performed in biological replicate with T-cells from 2 or 3 donors.

### Murine studies

Murine studies were conducted in accordance with UCSF Institutional Animal Care and Use Committee guidelines. NSG (NOD.Cg-*Prkdcscid Il2rgtm1Wjl*/SzJ, Jackson Laboratories) mice, 6-9 weeks old, male and female were bred at Preclinical Therapeutics Core at UCSF. Mice were injected intravenously via tail vein with 1e6 multiple MM tumor cells stably expressing luciferase. 6 days after tumor administration, tumor burden was quantified using bioluminescence and animals were randomized into treatment groups with equal average tumor burden. On day 7 after tumor injection mice were injected intravenously with 5e6 of CAR-T cells, Empty CAR or untransduced T cells at 1:1 ratio of CD4/CD8 T cells. Bioluminsecence imaging was done weekly to assess tumor burden (Perkin Elmer In Vivo Imaging System, Caliper Life Sciences). Survival end point of the study was determined by signs of symptomatic illness in the animals and required veterinary protocols for humane euthanasia.

### Murine blood CAR-T expansion assays

CAR-T and circulating tumor in vivo kinetics was analyzed every 2 weeks after CAR-Ts were injected in mice. Mouse blood was treated with 1X RBC lysis buffer (Biolegend 420301). White blood cells were stained with CD3, CD138 and CD70 for CAR-T (see table of antibodies) and tumor identification and analyzed by flow cytometry using a CytoFLEX Flow Cytometer (Beckman Coulter). Data was analyzed using FlowJo 10.10.0 software. CAR-T cells were identified by hCD3+ and GFP+ gate, cells per microliter count was calculated by multiplying total collected cell number by percent CAR-T cells and dividing by volume of mouse blood analyzed, 50 microliters.

### CAR-T cell in vitro immunophenotyping

CAR-T cells were cultured with tumor at 1:1 E:T ratio and after 24 hours exhaustion and memory phenotype was assessed by flow cytometry staining with CD3, CD45RA, CD62L, TIM3, LAG3 and PD1 markers (see table of antibodies for clone information). Cells were analyzed by flow cytometry and data was analyzed using FlowJo 10.10.0. Memory phenotype of CAR-T cells was determined using the following characteristics: stem cell memory (CD62L^+^CD45RA^+^), central memory (CD62L^+^CD45RA^-^), effector memory (CD62L^-^ CD45RA^-^) and effector T cells (CD62L^-^ CD45RA^+^).

### Quantitative flow cytometry for antigen density on cell lines

Cells were stained with CD70-APC and BCMA-APC flow cytometry antibody 1 ug/test (see table of antibodies for clone information). Number of CD70 and BCMA molecules per cell was determined using Quantum^TM^ APC MESF (Bangs Laboratories, 823) quantification beads, the beads were acquired with the same acquisition settings as cells. A set of beads is labeled with a standard number of APC fluorophores per bead to provide a calibration curve and equation to derive the number of antigen molecules per cell for the measured cell population. Using mean fluorescent intensity values and inputting them in the equation generated by standard curve, the number of molecules of CD70 and BCMA per cell was determined for each MM cell line.

### Cytokine analysis

CAR-Ts and untransduced T cells were cocultured with MM1S tumor cell line in triplicates at 1:1 ratio. The supernatant was collected, diluted 1:1 with RPMI media and snap frozen using liquid nitrogen. The analysis of cytokine levels was performed by Eve Technologies Corp. (Calgary, Alberta), using Luminex xMAP technology for multiplexed quantification. Multiplexing analysis was done with the Luminex™ 200 system (Luminex, Austin, TX, USA). Fourteen markers were simultaneously measured in the samples using Eve Technologies’ Human High Sensitivity 14-Plex Discovery Assay® (MilliporeSigma, Burlington, Massachusetts, USA) according to the manufacturer’s protocol. The markers tested included: GM-CSF, IFNγ, IL-1β, IL-2, IL-4, IL-5, IL-6, IL-8, IL-10, IL-12p70, IL-13, IL-17A, IL-23, TNF-α. Detection levels of these markers range from 0.11 – 3.25 pg/mL for the 14-plex. Each marker sensitivity values are available in the MilliporeSigma MILLIPLEX® MAP protocol.

### CD70 recombinant protein flow cytometry

Recombinant CD70 trimer protein conjugated to Alexa Fluor 647 (Acrobiosystems, cat. # CDL-HA246) was used to evaluate the affinity of truncated CD27-based CARs for their CD70 target. In a 96-well round-bottom plate, approximately 1105 CAR-T cells were resuspended in 50 μl of FACS buffer (DPBS + 2% FBS) containing 0.1 μg of CD70 recombinant protein, for a final concentration of 2 μg/ml. The cells were incubated at 4℃ for 1 hour, followed by washing twice in 200μl FACS buffer. Flow cytometry was performed on the CytoFLEX platform (Beckman Coulter), and data were analyzed using FlowJo_v10.7.2.

### ChIP-seq and ATAC-seq analysis

Raw data of ATAC-seq and H3K27Ac ChIP-seq data for MM patients and normal Memory B cells, Plasmablasts, and Plasma cells were downloaded from European Nucleotide Archive database with the Data Accession Number: PRJEB25605. Paired-end reads were mapped to the human reference genome (hg19) using Bowtie2 for ChIP-seq and ATAC-seq reads. Only reads mapping uniquely to the genome with not more than 2 mismatches were retained for further analysis. Clonal reads (i.e., reads mapping at the same genomic position and on the same strand) were collapsed into a single read. Peaks from ChIPseq or ATAC-seq were called using the MACS2 or Homer program. We also checked the potential motif binding in the CD70 promoter regions. We predicted the motif binding in the identified ATAC peak regions from MM patients using PROMO tool and ENCODE V3 336 and 161. We predicted the motif binding in the identified ATAC peak regions from MM patients using PROMO tool, and ENCODE V3 336 and 161

### Machine learning analysis for transcriptional regulation of CD70 expression

A list of 84 transcription factors was generated for this analysis by combining those from ATAC-seq motif analysis of MM patient tumors with analysis of ENCODE and PROMO data suggesting binding of different transcription factors to the *CD70* locus in memory B, plasmablast, plasma cells and MM cells. CD138+ enriched RNA-Seq data from 776 MM patients was incorporated into the model from CoMMpass-MMRF dataset (release IA19). All counts (in Transcripts per Million (TPM)) were log-transformed prior to model building and analysis. We developed an XGBoost model (Extreme Gradient Boosting) using the xgboost package in Python. The code used for our analysis is available at our repository here (https://github.com/BonellPatinoE/CD70-predictor-in-Myeloma.git) and the approach is similar to that we previously used for analysis of *CD38* and *CD46* regulators (*41, 55*) in MM except here applied to *CD70*. All transcription factors determined to be co-expressed were utilized together to build the model. We used randomized search with cross validation to find the best parameters for the XGBoost models to predict CD70 expression. Specifically we conducted a search over colsample_bytree, colsample_bylevel, colsample_bynode, gamma, learning_rate, max_depth, n_estimators, and subsample. Random combinations of these were used to build a model with 5-fold cross validation, for 5000 iterations, and the hyperparameters that gave the highest cross validation score was what was utilized in further model building. We conducted SHAP (Shapley Additive exPlanations) analysis on each model built to help describe what affect features have on *CD70* model predictions using the shap package. The SHAP analysis was conducted only with test data (20% of CoMMpass dataset). Model performance was evaluated by calculating coefficient of determination (R^2^) and adjusted R^2^ values. Furthermore, the model underwent 5-fold cross validation with the mean absolute error, R^2^, and adjusted R^2^ being reported for each fold to evaluate that performance was not dependent on our original choice of training and test data.

### CRISPR-Cas9 Gene Knockouts

Cas9 protein and sgRNA or scramble sgRNA were mixed in 1:2.5 molar ratio (Synthego Corporation) and incubated at 37 °C for 10–15 min. 1e6 of either primary CD3+ T cells or MM cell lines were spun down and washed with PBS, resuspended in 20 μL of P3 nucleofection/solution plus cas9/sgRNA mixture (P3 Primary Cell 4D-NucleofectorTM X Kit S) for primary T cells or SF Cell Line 4D-NucleofectorTM X Kit S (Lonza) for MM cell lines, and nucleofected using EO-115 or DS-137 nucleofection program in Lonza 4D-Nucleofector, respectively. 80 µL of warm (CTS™ OpTmizer™ T Cell Expansion SFM, Gibco™) or RPMI 1640 media (Gibco) was plated into each well and incubated at 37 °C for 15 min, then transferred to a 12-well plate supplemented with IL-7 and IL-15 for T-CD3+ T cells and 6-well plate for cell lines to recover. Then, based on the surface expression by flow cytometry, negative clones were sorted using FACSARIA-Fusion or FACSARIA III flow cytometer (BD Biosciences). sgRNA sequences were obtained from Brunello Library(*56*) as follows: *CD70*: CAGCTACGTATCCATCGTGA, AGCGCUGGATGCACACCACG; *TFAP2A*: AUCCUCGCAGGGACUACAGG; *CD27*: AGUGUGAUCCUUGCAUACCG. sgRNA sequence for *TNFRSF17* (BCMA) was obtained through from Synthego Corporation tool: GGUGUGACCAAUUCAGUGAA. Negative Control, scrambled sgRNA#1 and #2 mod-sgRNA specified by Synthego Corporation.

### Genomic DNA confirmation of CRISPR-Cas9 knockouts

DNA extraction from 1 million LP1 cells, whether fresh or frozen, was conducted using the Monarch® Genomic DNA Purification Kit (NEB, Cat # T3010S). Primers were procured from Integrated DNA Technologies and designed with the PrimerQuest™ Tool (IDT, Coralville, Iowa, USA) to yield a ~500 bp amplicon, ensuring a minimum of 200 bp upstream and downstream from the target sgRNA sequence. The primers were reconstituted in nuclease-free water to create a 10 µM stock. A PCR reaction employing 0.5 µM of both forward and reverse primers was carried out using the NEBNext® High-Fidelity 2X PCR Master Mix (Cat # M0541S) with genomic DNA ranging from 1 ng to 1 µg. Subsequently, the PCR product underwent purification using the QIAquick PCR Purification Kit (QIAGEN, Cat # 28104). Sanger Sequencing was outsourced to Quintara Biosciences (Fort Collins, CO, USA). The resulting Ab1 files for both control and experimental DNA groups were uploaded to the Synthego ICE analysis tool (Synthego Corporation, Redwood City, CA, USA; https://ice.synthego.com/#/), where indel %, Model Fit (R^2), and Knockout Score were employed to assess knockout efficiency. The primer sequences used for PCR amplification were as follows: TFAP2A: Set #1: Forward: CCACCCTACCAGCCTATCT, Reverse: ACTCTCCGCCTCCGAATAA. Set #2: Forward: CCCAATGCCGACTTCCA Reverse: CTGTATGTTCCAGGTATCCTTTCT. CD70: Set #1: Forward: CTGACAGGTTGAAGCAAGTAGA, Reverse: TCCCTGTTCTGGTCTCTGT. Set#2: Forward: GCAGTTTCCTTCCTTCCTTCTC, Reverse: CCTGTTCTGGTCTCTGTCTCT

### Immunoblot analysis

Nuclear lysates were prepared using the Nuclear Extracts Kit (Active Motif, # 40010). 20-50 ug Proteins were separated by SDS-PAGE and transferred into nitrocellulose membrane using the iBlot Transfer Stacks and the iBlot2 transfer system (ThermoFisher). Membranes were blocked using the Pierce™ Protein-Free Blocking Buffer (ThermoFisher). Primary antibodies targeting NSD2 (Abcam Cat# ab75359), HDAC2 (Millipore Cat# 05-814), H3 (Cell Signaling Technology Cat# 14269) and H3K36me2 (Abcam Cat# ab9049) were diluted in blocking buffer at 1:1000. Goat Anti-Rabbit Alexa Fluor Plus 680 (Thermo Fisher Scientific Cat# A32734, RRID:AB_2633283) and Goat Anti-Mouse Alexa Fluor Plus 800 (Thermo Fisher Scientific Cat# A32730, RRID:AB_2633279) secondary antibodies were used at dilution 1:10000. Membranes were scanned using an Li-Cor Odyssey CLx Imaging System (Li-Cor).

### Azacytidine Treatment of Cell Lines

AMO1 and MM1.S cell lines were treated with addition of freshly prepared azacytidine (5-Azacytidine, Selleckchem, Catalog No: S1782, Batch No. 07, resuspended in DMSO to make 1000X stocks) in fresh media every day of the treatment. Cells were either treated continuously for three days (3d) and allowed to recover for four days in fresh media (3d+4). On day 8, cells were assessed by flow cytometry, stained with CD70 antibodies and DRAQ7 viability dye (Biolegend, Catalog #: 424001).

### Mass spectrometry cell-surface proteomics

Data acquisition for AML and MM cell lines were performed using a timsTOF pro2 system. The acquisition of data-dependent acquisition (DDA) runs was carried out in PASEF mode within a mobility range of 0.7 to 1.3. The ramp time was set at 100 ms with a duty cycle of 100%, and the scan range was set from 100 m/z to 1700 m/z. The number of ramps varied based on gradient lengths to ensure sufficient data points for quantification, ranging between 4 and 10 ramps. Collision energy settings were optimized from 65 eV at 1/K0 of 1.6 Vs/cm2 to 20 at 1/K0 of 0.6 Vs/cm2, with a target intensity of 5,000 and intensity threshold of 1,000. MS2 scheduling was expedited, reducing the standard quadrupole switching time from 1.6 to 1.2 ms, and MS2 acquisition time was decreased from 2.75 to 2 ms. The raw LFQ spectral files were searched against human reviewed Uniprot database using LFQ-MBR workflow in Fragpipe 20.0 (MSFragger v4.0, IonQuant v1.10.12, Philosophor v5.1.0). In addition, trypsin and Lys-C were set as protease allowing up to two missed cleavages, methionine oxidation, and acetylation of N-termini were set as variable and carbamidomethylation of cysteine as fixed modification. For comparative analysis of CD70 expression in MM two independent datasets with primary patient data were downloaded from ProteomeXchange with ID PXD014165 and PXD022482. The raw spectra were analyzed using Fragpipe as mentioned above. Furthermore, the raw intensity for CD70 was log2 transformed, scaled, and normalized for comparative visualization of CD70 expression in various conditions.

## Supplementary Information

**Table S1 –.** Gene list from CoMMpass analysis (Excel file)

**Table S2.**
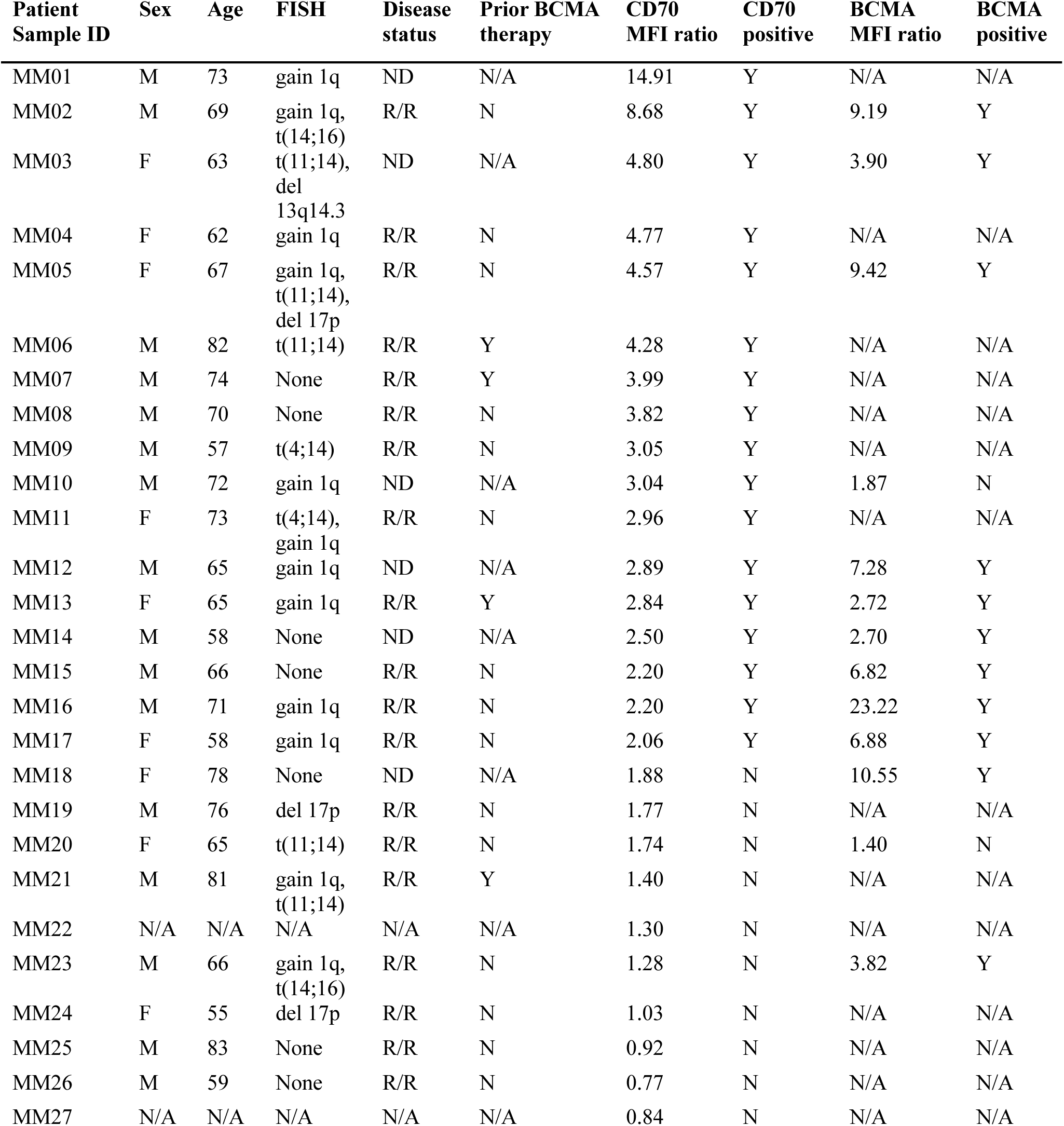
Patient characteristics for CD70 flow cytometry. Individual patient characteristics for primary patient samples analyzed using flow cytometry for CD70 and BCMA. Abbreviations used: M = male, F = female; ND = newly diagnosed; R/R = relapsed refractory; MFI = median fluorescent intensity; Y = yes, N= no, N/A = not available. MFI ratios are relative to negative (isotype) antibody control.

**Table S3 –.**
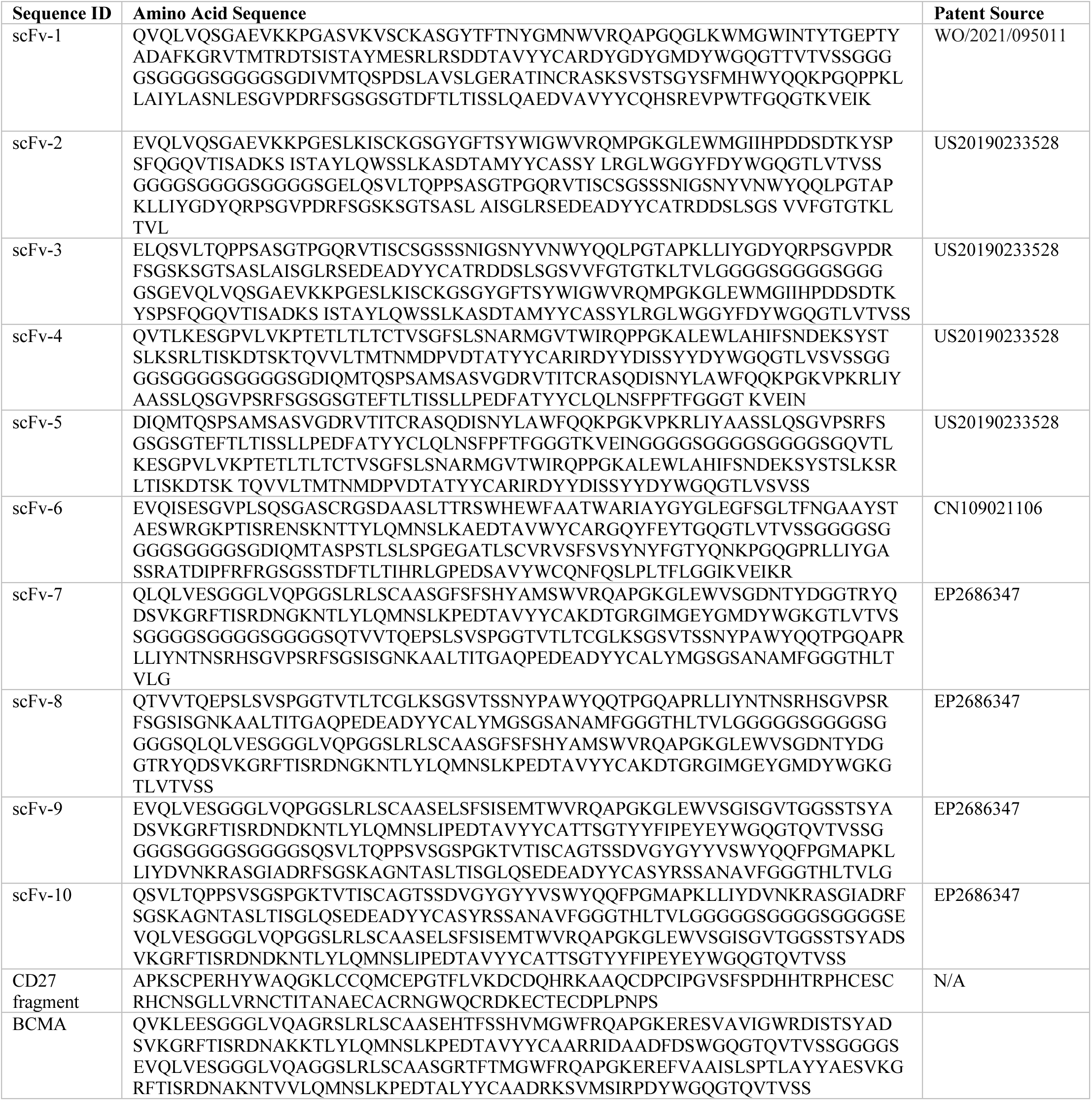
Sequences of scFv or other used in CAR-T designs. Amino acid sequences of tumor antigen-binding domain for each CAR-T design. For scFv-based CAR-Ts, the relevant source patent is shown at right.

**Table S4 –.**
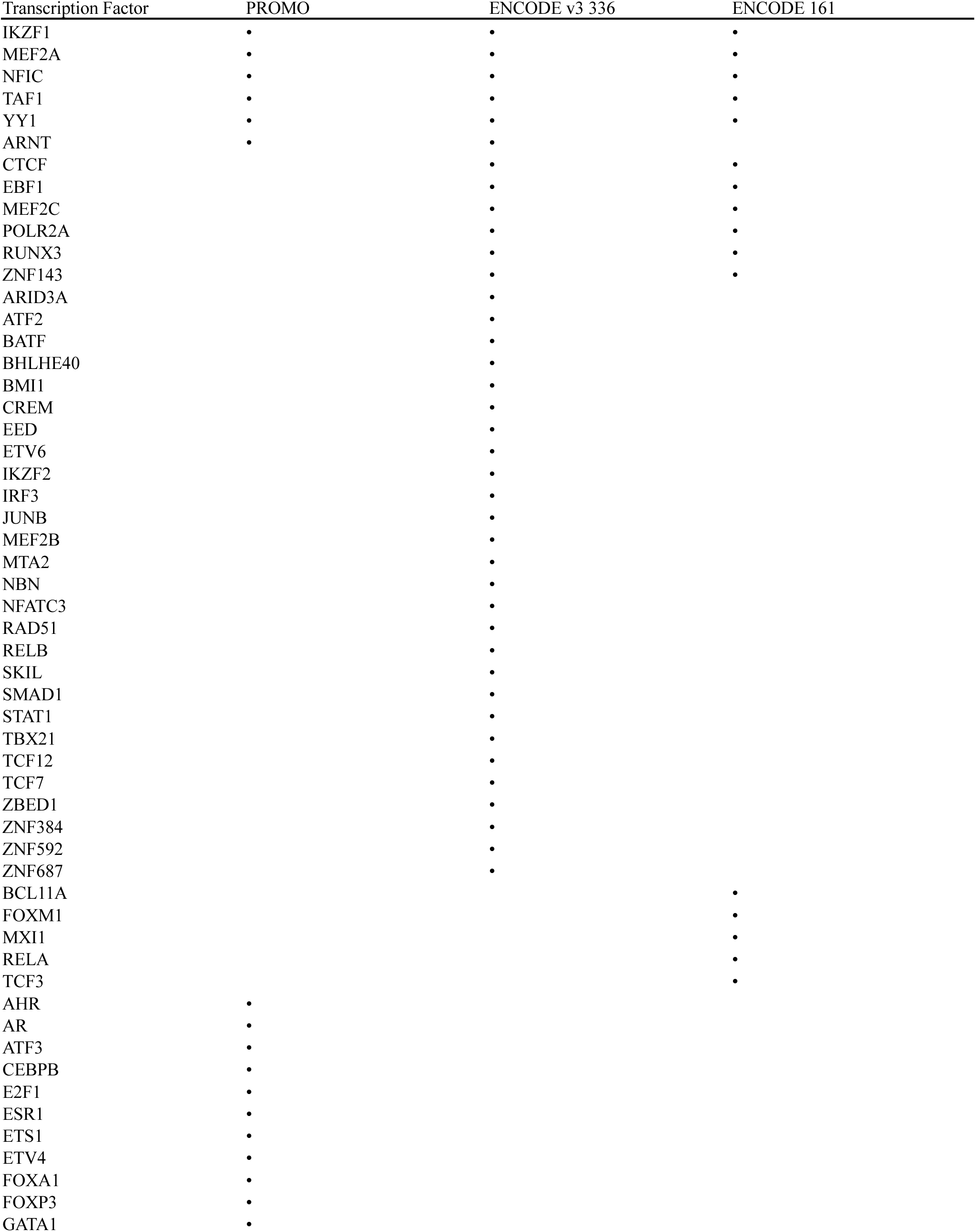
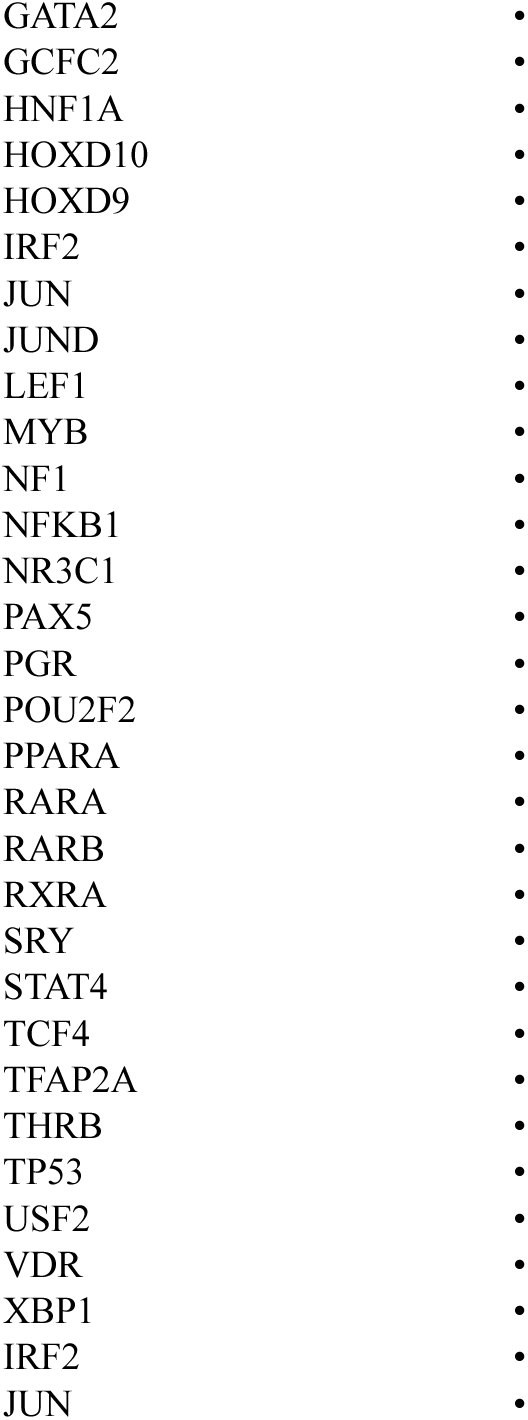
ChIP-seq and ATAC-seq peaks for epigenetic modeling. Transcription Factors predicted to bind CD70 promoter by PROMO or ENCODE. Potential transcription factors identified from ATAC-seq motif analysis on primary multiple myeloma patient samples or publicly available CHIP-Seq data.

| Transcription Factor | PROMO | ENCODE v3 336 | ENCODE 161 |
| --- | --- | --- | --- |
| IKZF1 | • | • | • |
| MEF2A | • | • | • |
| NFIC | • | • | • |
| TAF1 | • | • | • |
| YY1 | • | • | • |
| ARNT | • | • |  |
| CTCF |  | • | • |
| EBF1 |  | • | • |
| MEF2C |  | • | • |
| POLR2A |  | • | • |
| RUNX3 |  | • | • |
| ZNF143 |  | • | • |
| ARID3A |  | • |  |
| ATF2 |  | • |  |
| BATF |  | • |  |
| BHLHE40 |  | • |  |
| BMI1 |  | • |  |
| CREM |  | • |  |
| EED |  | • |  |
| ETV6 |  | • |  |
| IKZF2 |  | • |  |
| IRF3 |  | • |  |
| JUNB |  | • |  |
| MEF2B |  | • |  |
| MTA2 |  | • |  |
| NBN |  | • |  |
| NFATC3 |  | • |  |
| RAD51 |  | • |  |
| RELB |  | • |  |
| SKIL |  | • |  |
| SMAD1 |  | • |  |
| STAT1 |  | • |  |
| TBX21 |  | • |  |
| TCF12 |  | • |  |
| TCF7 |  | • |  |
| ZBED1 |  | • |  |
| ZNF384 |  | • |  |
| ZNF592 |  | • |  |
| ZNF687 |  | • |  |
| BCL11A |  |  | • |
| FOXM1 |  |  | • |
| MXI1 |  |  | • |
| RELA |  |  | • |
| TCF3 |  |  | • |
| AHR | • |  |  |
| AR | • |  |  |
| ATF3 | • |  |  |
| CEBPB | • |  |  |
| E2F1 | • |  |  |
| ESR1 | • |  |  |
| ETS1 | • |  |  |
| ETV4 | • |  |  |
| FOXA1 | • |  |  |
| FOXP3 | • |  |  |
| GATA1 | • |  |  |

|  |  |
| --- | --- |
| GATA2 | • |
| GCFC2 | • |
| HNF1A | • |
| HOXD10 | • |
| HOXD9 | • |
| IRF2 | • |
| JUN | • |
| JUND | • |
| LEF1 | • |
| MYB | • |
| NF1 | • |
| NFKB1 | • |
| NR3C1 | • |
| PAX5 | • |
| PGR | • |
| POU2F2 | • |
| PPARA | • |
| RARA | • |
| RARB | • |
| RXRA | • |
| SRY | • |
| STAT4 | • |
| TCF4 | • |
| TFAP2A | • |
| THRB | • |
| TP53 | • |
| USF2 | • |
| VDR | • |
| XBP1 | • |
| IRF2 | • |
| JUN | • |

**Figure S1.**
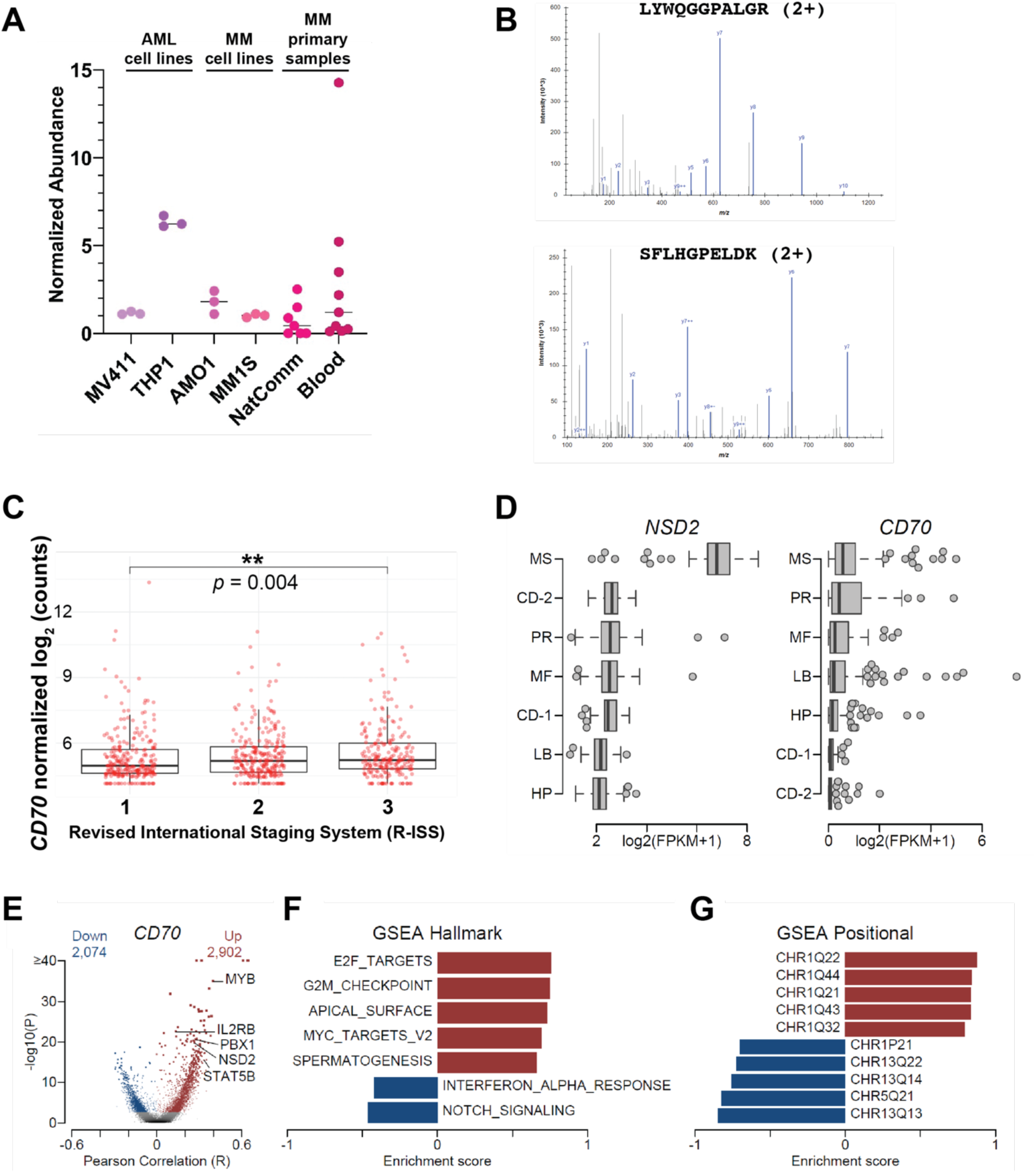
CD70 is expressed at the multiple myeloma surface. A. Cell surface capture proteomics of myeloma (AMO-1 and MM.1S) cell lines in comparison to acute myeloid leukemia (AML) cell lines indicated surface expression of CD70. Re-analysis of primary patient sample surface proteomic datasets (NatComm = Ferguson et al, Nature Communications (2022)(*1*); Blood = Anderson et al, Blood (2022)(*2*)) also indicates detectable CD70 in some samples. Plot shows normalized abundance based on mass spectrometric intensity. B. Example mass spectra from MM.1S cell line demonstrating peptides from CD70 identified by cell surface proteomics. C. CoMMpass transcript data (*n* = 776) demonstrating CD70 expression as a function of Revised International Staging System (R-ISS) stage (1, 2, 3) at diagnosis. D. Box plot of *NSD2* (left) and *CD70* (right) expression grouped by gene expression subtype in newly diagnosed multiple myeloma (NDMM) CoMMpass samples (*n* = 764). Gene expression subtypes are from Zhan et al. (CD-1: Cyclin D1; CD-2: Cyclin D1 and CD20; HP: Hyperdiploid; LB: Low Bone Disease; MF: MAF; MS: MMSET; PR: Proliferation) E. Volcano plot of genes associated with *CD70* expression in CoMMpass NDMM samples (*n* = 764). Genes significantly associated with *CD70* (FDR <0.01) are shown in red (positive) and blue (negative) with the numbers listed above and select genes labelled. F. Top Gene Set Enrichment Analysis (GSEA) Hallmark pathways associated with *CD70* expression. Top 5 pathways positively (red) and negatively (blue) associated with *CD70* expression are shown, and only pathways with an FDR < 0.01 are included. G. Top GSEA positional gene sets associated with *CD70* expression. Only the top 5 pathways positively and negatively associated with *CD70* expression are shown.

**Figure S2.**
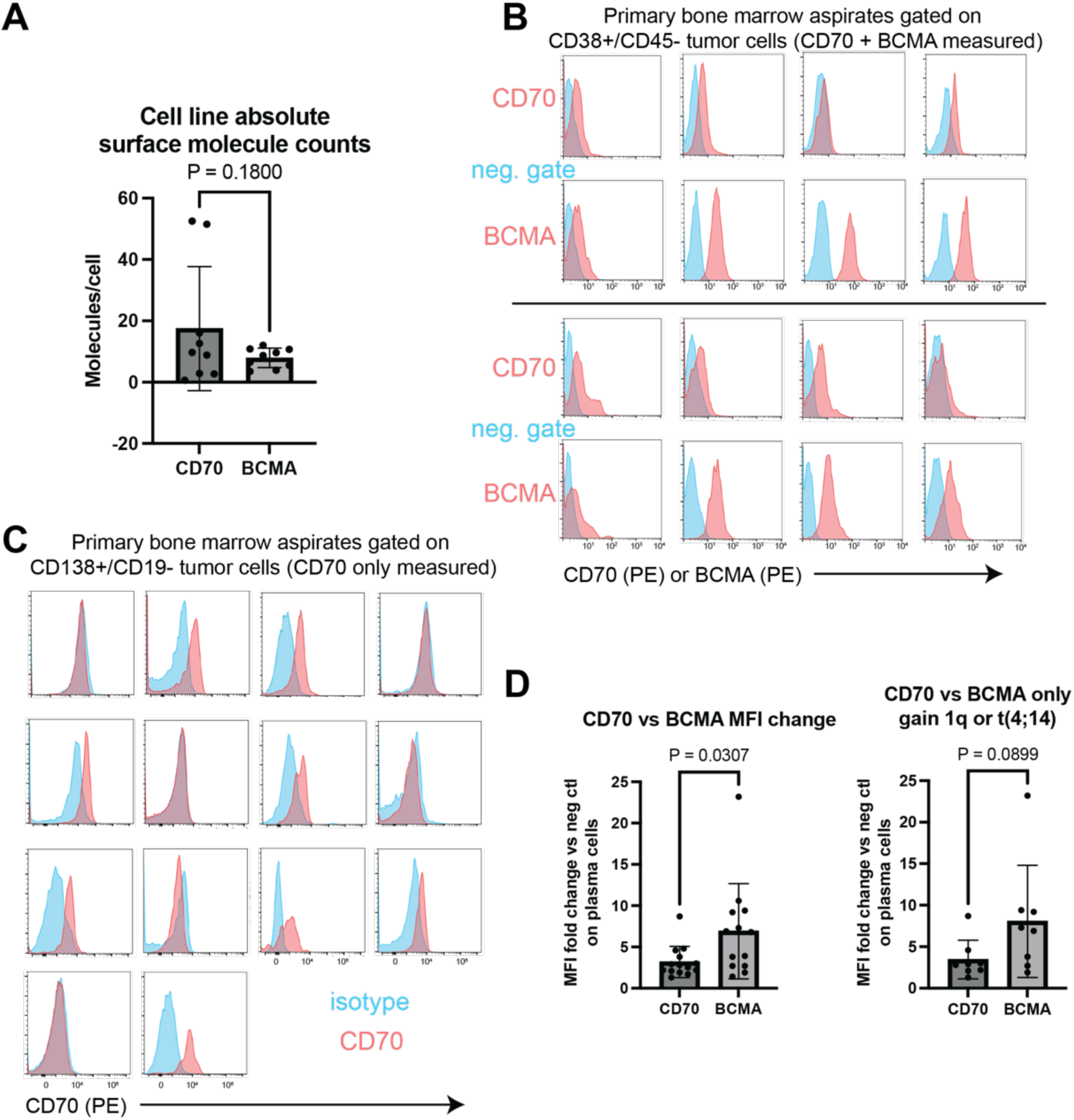
CD70 and BCMA expression on myeloma models and patient samples. A. Absolute quantification of surface molecules of CD70 and BCMA on myeloma cell lines (displayed in Fig. 1D) shows non-significant difference between CD70 and BCMA. *n* = 8 cell lines, performed in biological duplicate. *p*-value by two-tailed *t*-test. B. Flow cytometry of primary patient sample bone marrow aspirate with measurement of both CD70 and BCMA on CD38+/CD45dim/− mononuclear cell population. Signal shown (APC fluorophore for both antibodies) compared to unstained control. Each sample tested once. C. Flow cytometry of primary patient sample bone marrow aspirate with measurement of CD70 (only) on CD138+/CD19− mononuclear cells. Each sample tested once. D. For samples with measurement of both CD70 and BCMA, comparison of mean fluorescence intensity between patient plasma cells and negative controls. (left) For all samples (*n* = 13), BCMA shows moderately increased intensity compared to CD70. (right) For 8 samples with FISH results showing gain of Chr1q or t(4;14), BCMA increase was non-significant compared to CD70. *p*-value by two tailed *t*-test.

**Figure S3.**
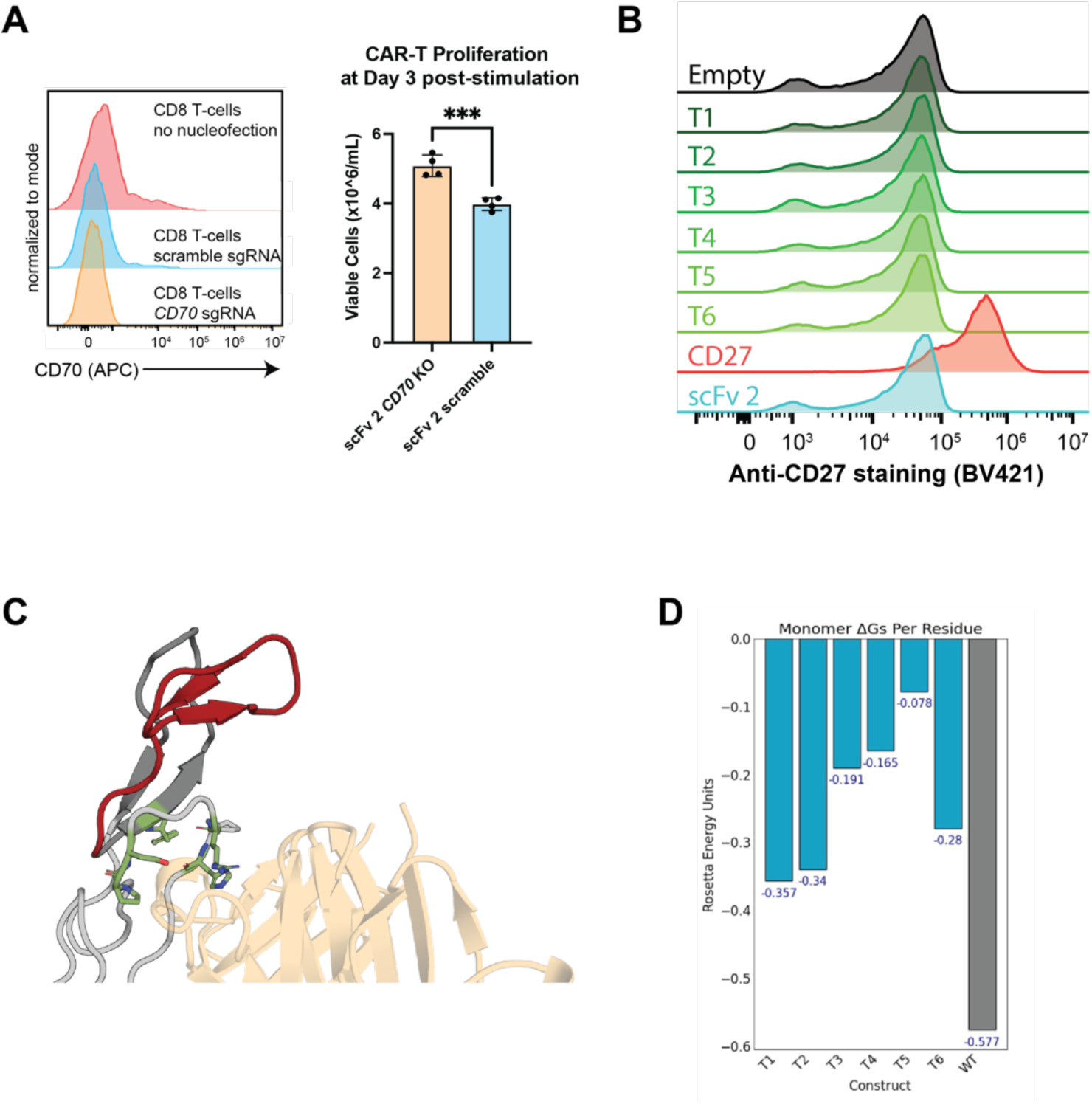
Further investigation of the CD70:CD27 interaction including Rosetta analysis. A. Flow cytometry on donor T-cells after nucleofection with recombinant Cas9 complexed with *CD70*-targeting sgRNA. B. Flow cytometry of CD70 KO donor T-cells transduced with the noted CD27 CAR truncation variants (see Fig. 2 of main text), stained with anti-CD27 monoclonal antibody. Only the full-length extracellular domain construct (“CD27”) showed any discernible staining. C. Portion of the CD27/CD70 complex structure with CD70 colored in tan, CRD1 colored in dark gray with truncated portions in construct T1 colored in red, and CRD2 colored in light gray. Interface positions in the non-truncated part of CRD1 and CRD2 that are predicted by Rosetta to contribute substantially to binding affinity with CD70 (hot spots, defined as change in binding energy upon mutation ΔΔG > 1 Rosetta Energy Unit) are shown in green stick representation. D. Rosetta-predicted stability (ΔG) scores, normalized by total residue number, for each CD27 CRD1 truncation construct as a monomer (see Supplementary Methods for additional details).

**Figure S4.**
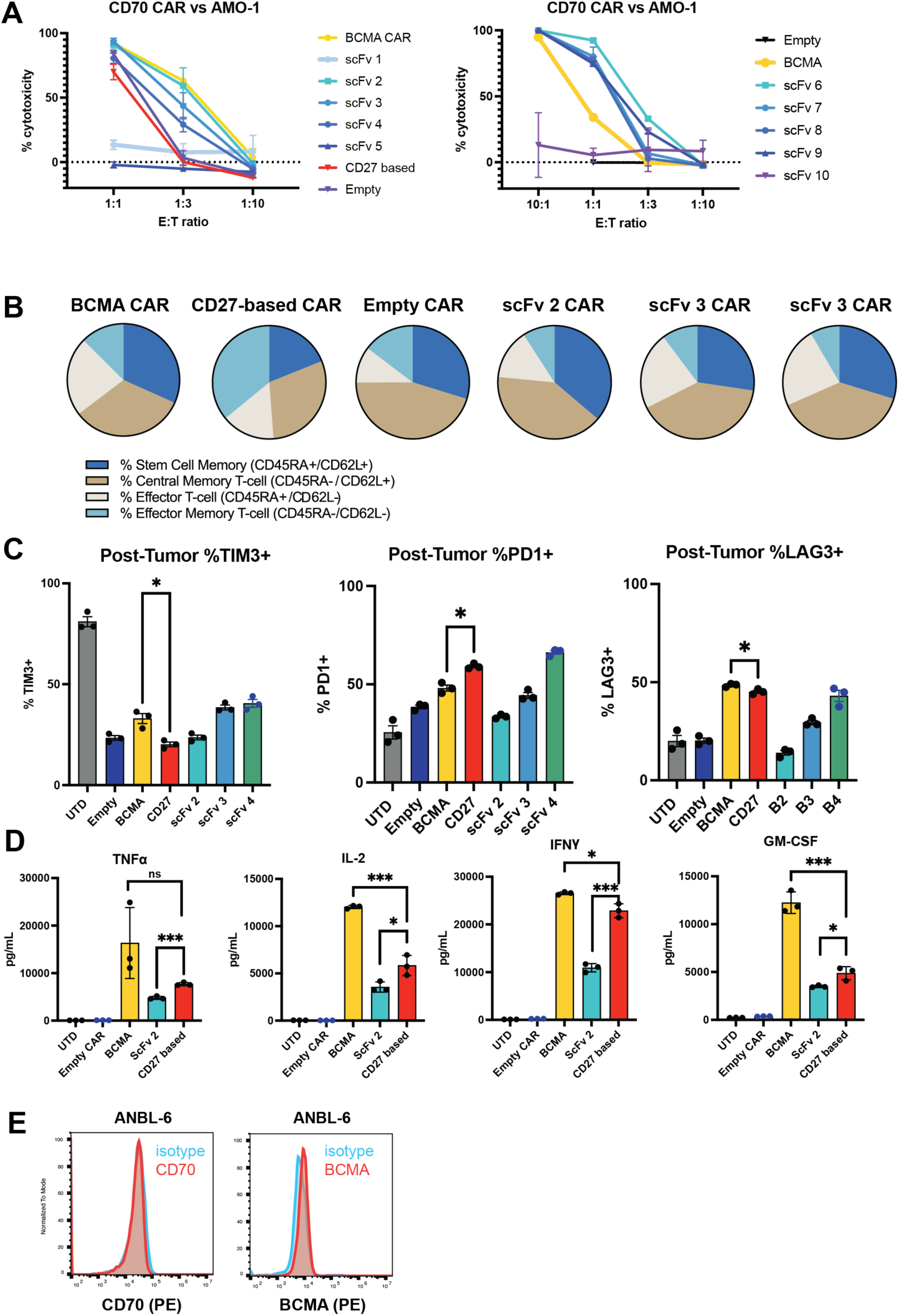
In vitro characterization of anti-CD70 CAR-Ts. A. In vitro screening of 10 scFv-based CAR designs versus AMO-1 myeloma cells, compared to BCMA CAR and FL CD27 ECD-based CAR. 18 hr co-culture at noted effector:tumor (E:T) ratio. *n* = 4 technical replicates. B. Flow cytometric analysis of indicated CAR constructs after 24 hr co-culture with LP-1 cells (1:1 E:T) to assess stem-ness properties, as measured by positivity for CD62L and CD45RA staining. Pie charts demonstrate percentage of cells falling into each phenotypic state (stem cell memory (CD62L^+^CD45RA^+^), central memory (CD62L^+^CD45RA^-^), effector memory (CD62L^-^ CD45RA^-^) and effector T cells (CD62L^-^ CD45RA^+^)). *n* = 3 technical replicates. C. Flow cytometry of same study as in B. with staining for PD-1, LAG-3, and TIM-3 to assess activation and exhaustion state after LP-1 co-culture at 1:1 E:T, 24 hrs. *n* = 3 technical replicates. D. Quantification of specific individual cytokines from heatmaps shown in Fig. 3B. Performed in biological triplicate after co-culture with indicated CARs and LP-1 cells. *p*-value by two-tailed *t*-test. \**p* <0.05, \*\**p* < 0.01, \*\*\*\**p* < 0.005. E. Flow cytometric analysis of ANBL-6 cell line demonstrating no detectable CD70 and very low BCMA. Depicted is one of 2 biological replicates.

**Figure S5.**
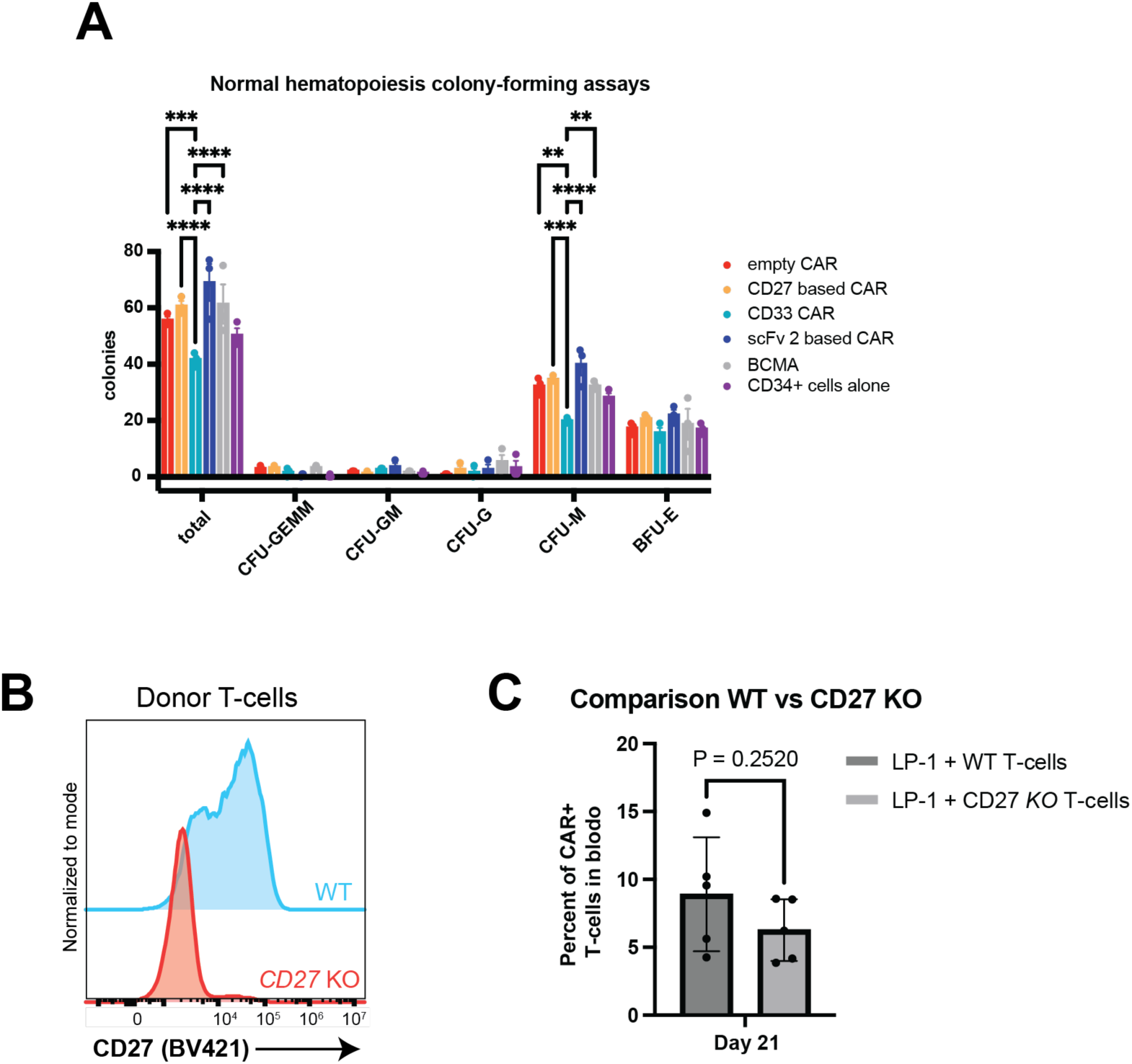
Probing the CD27-based CAR in vivo expansion phenotype. A. Colony-forming clonogenic assays (see Supplementary Methods) show no difference in hematopoietic precursor formation after treatment with CD70-targeting CAR-Ts. anti-CD33 CAR-T(*3*) used as positive control. B. Flow cytometric analysis of donor T-cells confirming successful knockout of endogenous CD27 via Cas9 ribonucleoprotein prior to CAR transduction. Presented is one of two biological replicates. C. Comparison of CD27 CAR-T cell expansion in murine peripheral blood at Day 21 of study, with WT CAR T-cells or *CD27* KO CAR T-cells. CD27 KO did not lead to a significant decrease in CAR expansion. Data from *n* = 5 mice per arm. *p*-value by two-tailed *t*-test.

**Figure S6.**
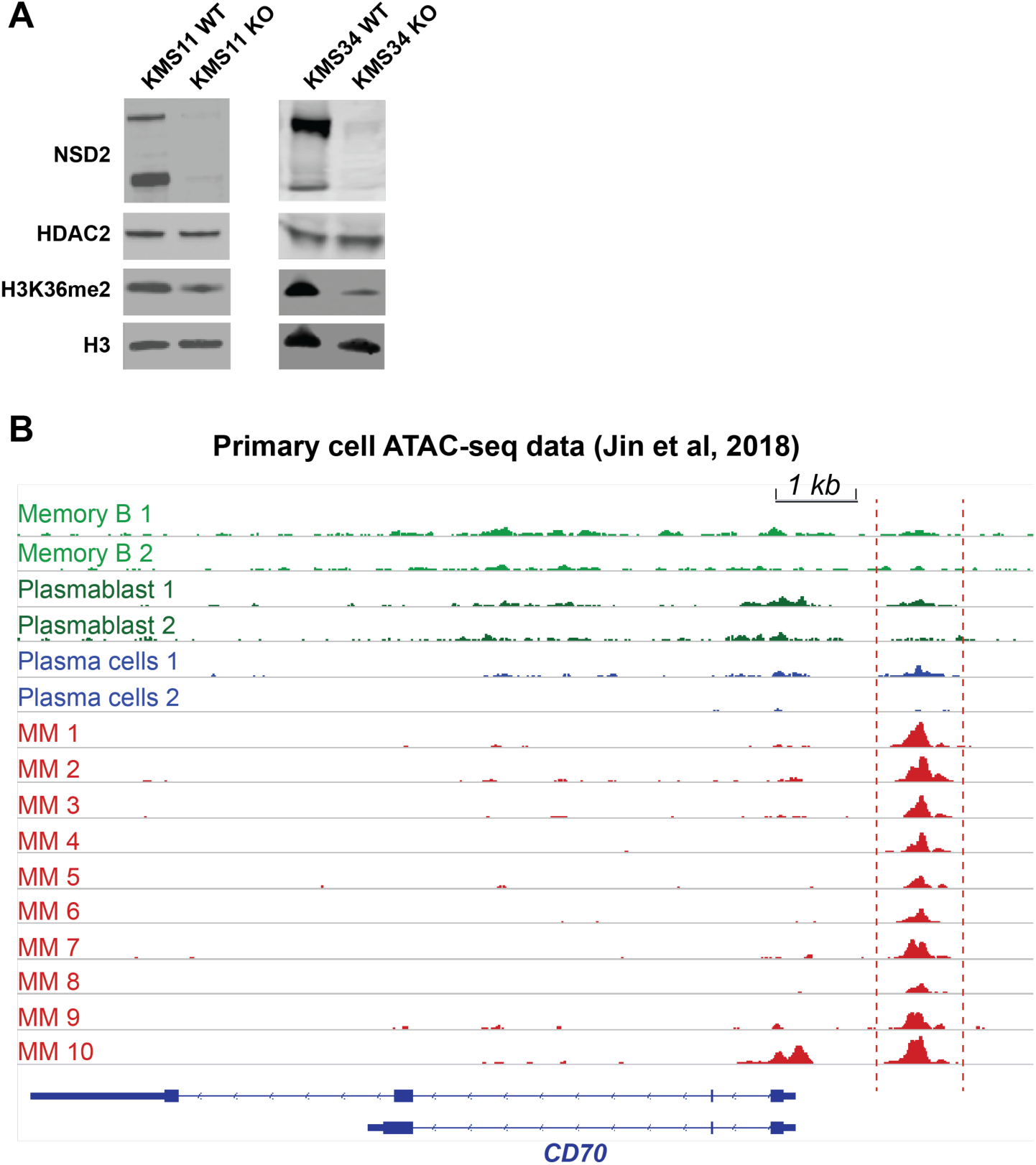
Regulation of CD70 in myeloma. A. Immunoblots confirming NSD2 knockout in KMS11 and KMS34 cell lines. H3K36me2 signal decrease confirms functional loss of histone methylation in the context of NSD2 loss. Total HDAC2 and Histone H3 levels are unchanged. B. Called peaks from ATAC-seq (data from Jin et al (2018)(*4*) at the *CD70* locus, across normal memory B-cells (MB), plasmablasts, normal plasma cells, and malignant plasma cells derived from multiple myeloma (MM) patients (ten illustrated here; 24 total included in study). Dashed red lines highlight major ATAC-seq peak present only in myeloma patient samples. Transcription factor motifs (listed in Table S4) extracted from this peak.

**Figure S7.**
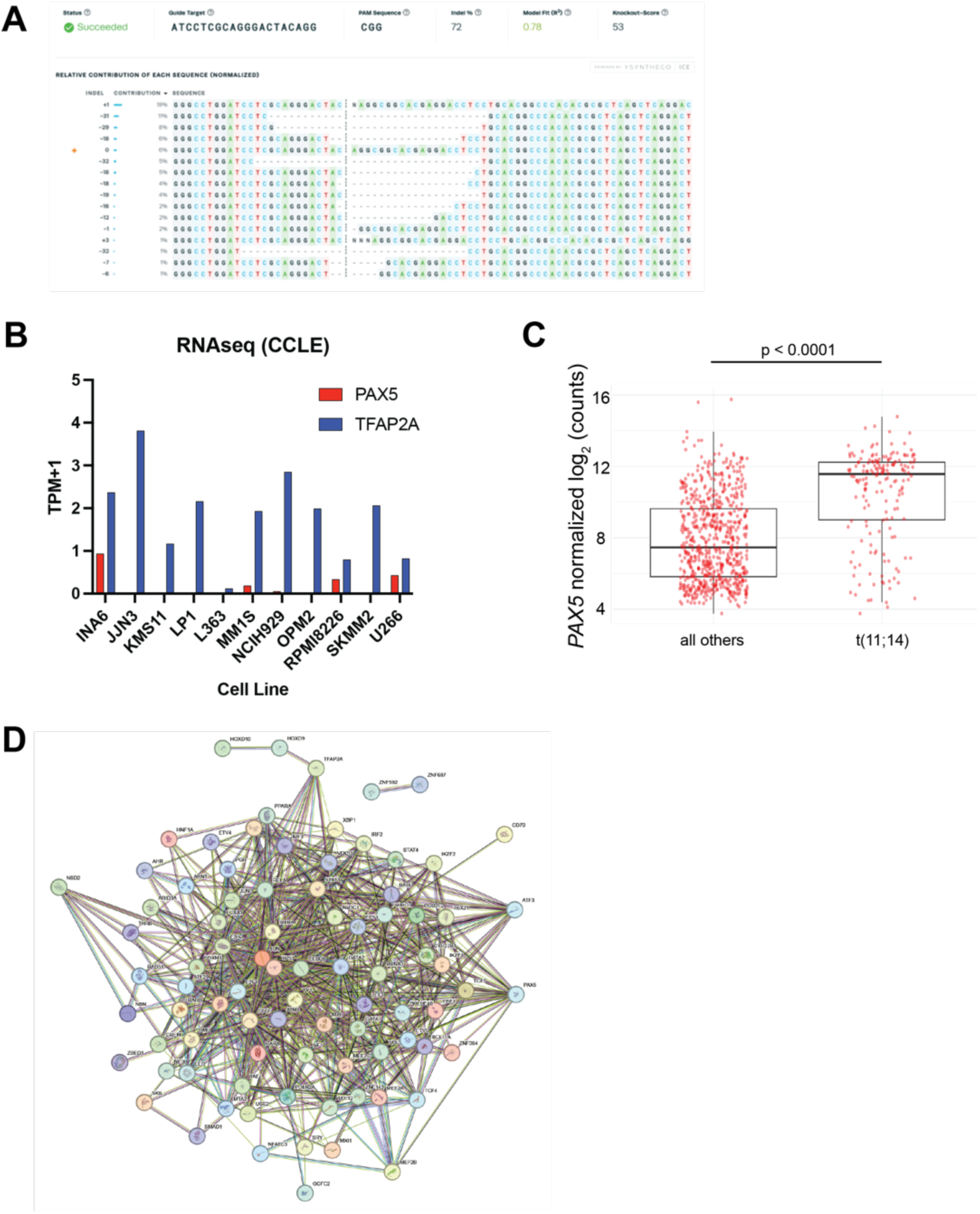
Interrogating transcription factors regulating *CD70*. A. Synthego analysis tool demonstrates successful knockout of *TFAP2A* in bulk LP-1 cells, with 72% knockout based on genomic DNA analysis by Sanger sequencing B. Transcript levels of *TFAP2A* and *PAX5* in the most commonly-used MM cell lines (from Sarin et al (2020)(*5*)) demonstrates minimal expression of *PAX5*. Bulk RNA-seq data from Cancer Cell Line Encyclopedia. C. Transcipt analysis of the CoMMpass database (n = 776) demonstrates elevated *PAX5* expression in t(11;14) myeloma compared to other genotypes. *p*-value by two-tailed *t*-test. D. STRING analysis of all transcription factors predicted to regulate CD70 based on ChIP-seq and ATAC-seq analysis (Table S4) shows strong network effects with many mutually interacting proteins.

**Figure S8.**
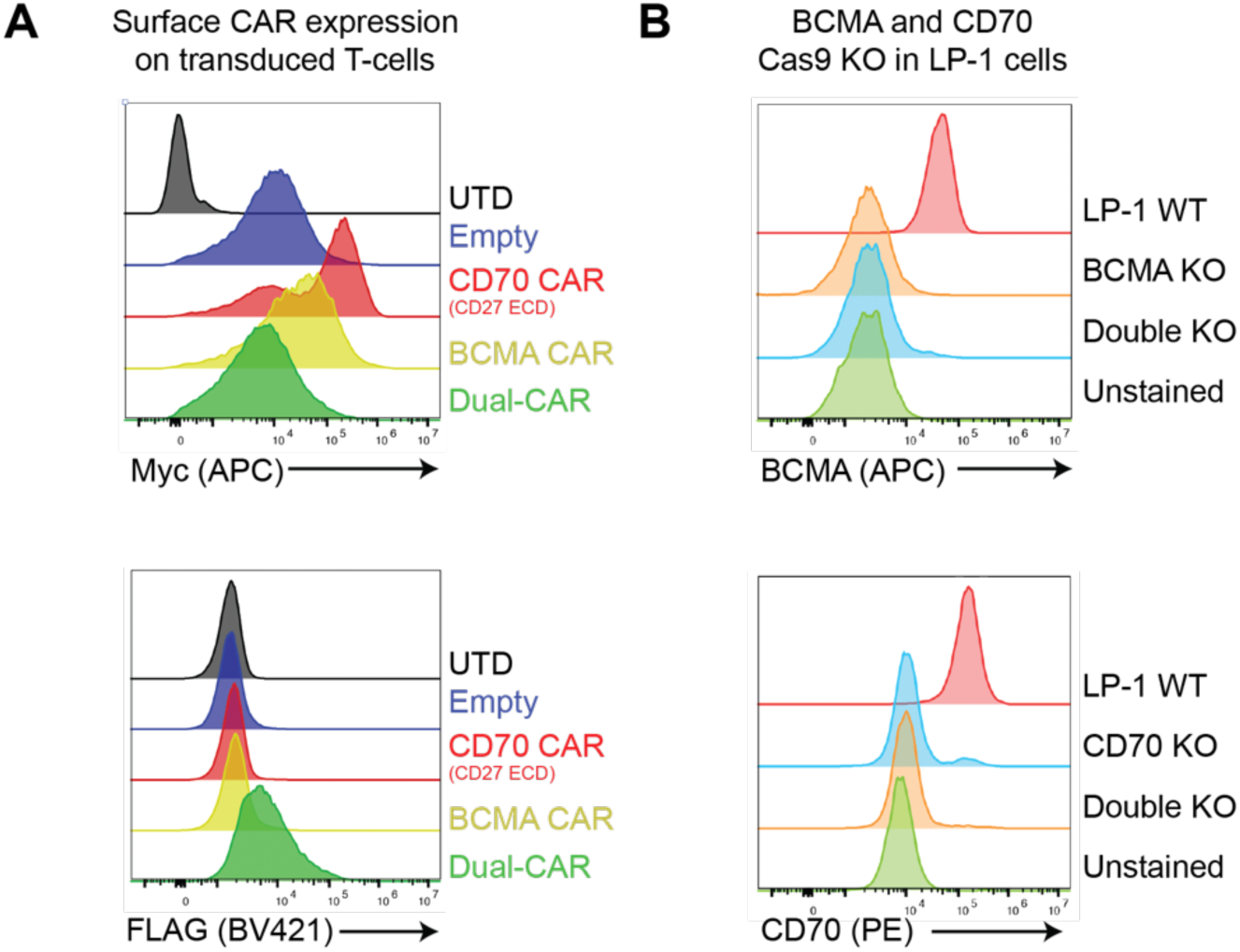
Dual-targeting CAR T cells against BCMA and CD70. A. (*top*) Flow cytometric analysis confirming successful expression of all single targeting CAR constructs as well as CD70-targeting CAR (FL CD27 ECD-based) in dual-targeting CAR based on myc tag staining at N-terminus of the CAR. (*bottom*) Flow cytometry confirms expression of the BCMA-targeting CAR (derived from cilta-cel biparatopic VHH sequence) only present in dual-targeting CAR based on FLAG tag (see CAR designs in **Fig. 7B**). UTD = untransduced. Representative of *n* = 2 biological replicates. B. Generation of LP-1 myeloma knockout cell lines. LP-1 nucleofected with Cas9 RNP complexed with sgRNA targeting *TNFSRF17A* (encoding BCMA) or sgRNA targeting *CD70*, in each single knockout cell lines, or simultaneous nucleofection for “Double KO”. In bulk cell population, flow cytometry confirms successful knockout of each gene in CD70 KO and BCMA KO lines, respectively, and knockout of both genes in Double KO cell lines.

### Supplementary Methods

#### Clonogenic assay

1,000 CD34+ cells from healthy donor mobilized peripheral blood were co-incubated with 1,000 CAR-T, empty CAR-T cells, or medium only (IMDM, 2% FBS) at an E:T ratio of 1:1 for 5 hours in V-bottom 96 well plates. Cells were then plated in triplicate in methylcellulose-based medium supplemented with recombinant cytokines (MethoCult H4434 Classic, STEMCELL Technologies). After 13-14 days, colonies were classified and counted as CFU-GEMM, -GM, -G, -M, or BFU-E.

#### Rosetta Analysis

PDB 7KX0 chains F, A, and B were used for all analyses. To prepare structure for Rosetta analyses, we removed all heteroatoms and minimized the structure using a constrained Rosetta Relax protocol using the script described in the Supplement of Cao et al. (*6*). To calculate ΔG values of the monomer, truncated chains were isolated from the relaxed trimer structure and scored individually using the Rosetta REF2015 scorefunction. Interface binding hotspots were identified using the Rosetta Flex ddG protocol (*7*) to mutate interface positions to Alanines and determine resultant ΔΔG values for the mutated complex. Mutated interfaces that yielded a ΔΔG value > 1 (subtracting the estimated binding energy of the wild-type complex from the mutated complex) were designated as interface binding hotspots.

## References

1. S. A. Holstein, S. J. Grant, T. M. Wildes, Chimeric Antigen Receptor T-Cell and Bispecific Antibody Therapy in Multiple Myeloma: Moving Into the Future. J Clin Oncol 41, 4416–4429 (2023).

2. R. S. Firestone, S. Mailankody, Current use of CAR T cells to treat multiple myeloma. Hematology Am Soc Hematol Educ Program 2023, 340–347 (2023).

3. D. K. Hansen et al., Idecabtagene Vicleucel for Relapsed/Refractory Multiple Myeloma: Real-World Experience From the Myeloma CAR T Consortium. J Clin Oncol 41, 2087–2097 (2023).

4. T. Martin et al., Ciltacabtagene Autoleucel, an Anti-B-cell Maturation Antigen Chimeric Antigen Receptor T-Cell Therapy, for Relapsed/Refractory Multiple Myeloma: CARTITUDE-1 2-Year Follow-Up. J Clin Oncol 41, 1265–1274 (2023).

5. O. Van Oekelen et al., Interventions and outcomes of patients with multiple myeloma receiving salvage therapy after BCMA-directed CAR T therapy. Blood 141, 756–765 (2023).

6. A. D. Cohen et al., B cell maturation antigen-specific CAR T cells are clinically active in multiple myeloma. J Clin Invest 129, 2210–2221 (2019).

7. H. Lee et al., Mechanisms of antigen escape from BCMA- or GPRC5D-targeted immunotherapies in multiple myeloma. Nat Med 29, 2295–2306 (2023).

8. E. L. Smith et al., GPRC5D is a target for the immunotherapy of multiple myeloma with rationally designed CAR T cells. Sci Transl Med 11, (2019).

9. S. Mailankody et al., GPRC5D-Targeted CAR T Cells for Myeloma. N Engl J Med 387, 1196–1206 (2022).

10. A. K. Mishra et al., CAR-T-Cell Therapy in Multiple Myeloma: B-Cell Maturation Antigen (BCMA) and Beyond. Vaccines (Basel) 11, (2023).

11. M. Uhlen et al., A genome-wide transcriptomic analysis of protein-coding genes in human blood cells. Science 366, (2019).

12. J. Jacobs et al., CD70: An emerging target in cancer immunotherapy. Pharmacol Ther 155, 1–10 (2015).

13. T. Flieswasser et al., Screening a Broad Range of Solid and Haematological Tumour Types for CD70 Expression Using a Uniform IHC Methodology as Potential Patient Stratification Method. Cancers (Basel) 11, (2019).

14. T. Flieswasser et al., The CD70-CD27 axis in oncology: the new kids on the block. J Exp Clin Cancer Res 41, 12 (2022).

15. R. Q. Hintzen et al., CD70 represents the human ligand for CD27. Int Immunol 6, 477–480 (1994).

16. C. Riether et al., Targeting CD70 with cusatuzumab eliminates acute myeloid leukemia stem cells in patients treated with hypomethylating agents. Nat Med 26, 1459–1467 (2020).

17. F. Perna et al., Integrating Proteomics and Transcriptomics for Systematic Combinatorial Chimeric Antigen Receptor Therapy of AML. Cancer Cell 32, 506–519 e505 (2017).

18. J. A. McEarchern et al., Preclinical characterization of SGN-70, a humanized antibody directed against CD70. Clin Cancer Res 14, 7763–7772 (2008).

19. S. H. Panowski et al., Preclinical Development and Evaluation of Allogeneic CAR T Cells Targeting CD70 for the Treatment of Renal Cell Carcinoma. Cancer Res 82, 2610–2624 (2022).

20. M. B. Nilsson et al., CD70 is a therapeutic target upregulated in EMT-associated EGFR tyrosine kinase inhibitor resistance. Cancer Cell 41, 340–355 e346 (2023).

21. C. Riether et al., Tyrosine kinase inhibitor-induced CD70 expression mediates drug resistance in leukemia stem cells by activating Wnt signaling. Sci Transl Med 7, 298ra119 (2015).

22. M. B. Leick et al., Non-cleavable hinge enhances avidity and expansion of CAR-T cells for acute myeloid leukemia. Cancer Cell 40, 494–508 e495 (2022).

23. T. Sauer et al., CD70-specific CAR T cells have potent activity against acute myeloid leukemia without HSC toxicity. Blood 138, 318–330 (2021).

24. S. Srour et al., paper presented at the Cancer Res (Supplement), Orlando, FL, 2023.

25. I. D. Ferguson et al., The surfaceome of multiple myeloma cells suggests potential immunotherapeutic strategies and protein markers of drug resistance. Nat Commun 13, 4121 (2022).

26. G. S. F. Anderson et al., Unbiased cell surface proteomics identifies SEMA4A as an effective immunotherapy target for myeloma. Blood 139, 2471–2482 (2022).

27. A. Palumbo et al., Revised International Staging System for Multiple Myeloma: A Report From International Myeloma Working Group. J Clin Oncol 33, 2863–2869 (2015).

28. A. Schavgoulidze et al., Biallelic deletion of 1p32 defines ultra-high-risk myeloma, but monoallelic del(1p32) remains a strong prognostic factor. Blood 141, 1308–1315 (2023).

29. F. Zhan et al., The molecular classification of multiple myeloma. Blood 108, 2020–2028 (2006).

30. D. R. Shaffer et al., T cells redirected against CD70 for the immunotherapy of CD70-positive malignancies. Blood 117, 4304–4314 (2011).

31. X. Wang et al., The proteasome deubiquitinase inhibitor VLX1570 shows selectivity for ubiquitin-specific protease-14 and induces apoptosis of multiple myeloma cells. Sci Rep 6, 26979 (2016).

32. M. A. Nix, A. P. Wiita, Alternative target recognition elements for chimeric antigen receptor (CAR) T cells: beyond standard antibody fragments. Cytotherapy In press, (2024).

33. Q. J. Wang et al., Preclinical Evaluation of Chimeric Antigen Receptors Targeting CD70-Expressing Cancers. Clin Cancer Res 23, 2267–2276 (2017).

34. W. Liu et al., Structural delineation and phase-dependent activation of the costimulatory CD27:CD70 complex. J Biol Chem 297, 101102 (2021).

35. J. Cheng et al., Revealing the impact of CD70 expression on the manufacture and functions of CAR-70 T-cells based on single-cell transcriptomics. Cancer Immunol Immunother 72, 3163–3174 (2023).

36. J. K. Leman et al., Macromolecular modeling and design in Rosetta: recent methods and frameworks. Nat Methods 17, 665–680 (2020).

37. M. V. Maus, M. B. Leick. (USA, 2022).

38. J. A. Fraietta et al., Determinants of response and resistance to CD19 chimeric antigen receptor (CAR) T cell therapy of chronic lymphocytic leukemia. Nat Med 24, 563–571 (2018).

39. D. L. Porter et al., Chimeric antigen receptor T cells persist and induce sustained remissions in relapsed refractory chronic lymphocytic leukemia. Sci Transl Med 7, 303ra139 (2015).

40. R. Popovic et al., Histone methyltransferase MMSET/NSD2 alters EZH2 binding and reprograms the myeloma epigenome through global and focal changes in H3K36 and H3K27 methylation. PLoS Genet 10, e1004566 (2014).

41. P. Choudhry et al., Functional multi-omics reveals genetic and pharmacologic regulation of surface CD38 in multiple myeloma. BioRxiv (preprint), (2021).

42. Y. Jin et al., Active enhancer and chromatin accessibility landscapes chart the regulatory network of primary multiple myeloma. Blood 131, 2138–2150 (2018).

43. C. Cobaleda, A. Schebesta, A. Delogu, M. Busslinger, Pax5: the guardian of B cell identity and function. Nat Immunol 8, 463–470 (2007).

44. V. A. Gupta et al., Venetoclax sensitivity in multiple myeloma is associated with B-cell gene expression. Blood 137, 3604–3615 (2021).

45. S. Ghorashian et al., CD19/CD22 targeting with cotransduced CAR T cells to prevent antigen-negative relapse after CAR T-cell therapy for B-cell ALL. Blood 143, 118–123 (2024).

46. C. Fernandez de Larrea et al., Defining an Optimal Dual-Targeted CAR T-cell Therapy Approach Simultaneously Targeting BCMA and GPRC5D to Prevent BCMA Escape-Driven Relapse in Multiple Myeloma. Blood Cancer Discov 1, 146–154 (2020).

47. P. Rodriguez-Otero et al., GPRC5D as a novel target for the treatment of multiple myeloma: a narrative review. Blood Cancer J 14, 24 (2024).

48. O. Landgren, D. Kazandjian, Modern Myeloma Therapy + Sustained Minimal Residual Disease-Negative = (Functional) Cure! J Clin Oncol 40, 2863–2866 (2022).

49. Y. Wang et al., Humoral immune reconstitution after anti-BCMA CAR T-cell therapy in relapsed/refractory multiple myeloma. Blood Adv 5, 5290–5299 (2021).

50. D. Bausch-Fluck et al., The in silico human surfaceome. Proc Natl Acad Sci U S A 115, E10988–E10997 (2018).

51. M. I. Love, W. Huber, S. Anders, Moderated estimation of fold change and dispersion for RNA-seq data with DESeq2. Genome Biol 15, 550 (2014).

52. A. Zhu, J. G. Ibrahim, M. I. Love, Heavy-tailed prior distributions for sequence count data: removing the noise and preserving large differences. Bioinformatics 35, 2084–2092 (2019).

53. L. Kolberg, U. Raudvere, I. Kuzmin, J. Vilo, H. Peterson, gprofiler2 -- an R package for gene list functional enrichment analysis and namespace conversion toolset g:Profiler. F1000Res 9, (2020).

54. M. Hudecek et al., The nonsignaling extracellular spacer domain of chimeric antigen receptors is decisive for in vivo antitumor activity. Cancer Immunol Res 3, 125–135 (2015).

55. A. Wadhwa et al., CD46-targeted theranostics for Positron Emission Tomography and 225Ac-Radiopharmaceutical Therapy of Multiple Myeloma. Clin Cancer Res, (2023).

56. J. G. Doench et al., Optimized sgRNA design to maximize activity and minimize off-target effects of CRISPR-Cas9. Nat Biotechnol 34, 184–191 (2016).

## Supplementary References

1. I. D. Ferguson et al., The surfaceome of multiple myeloma cells suggests potential immunotherapeutic strategies and protein markers of drug resistance. Nat Commun 13, 4121 (2022).

2. G. S. F. Anderson et al., Unbiased cell surface proteomics identifies SEMA4A as an effective immunotherapy target for myeloma. Blood 139, 2471–2482 (2022).

3. K. Mandal et al., Structural surfaceomics reveals an AML-specific conformation of integrin beta(2) as a CAR T cellular therapy target. Nat Cancer 4, 1592–1609 (2023).

4. Y. Jin et al., Active enhancer and chromatin accessibility landscapes chart the regulatory network of primary multiple myeloma. Blood 131, 2138–2150 (2018).

5. V. Sarin et al., Evaluating the efficacy of multiple myeloma cell lines as models for patient tumors via transcriptomic correlation analysis. Leukemia 34, 2754–2765 (2020).

6. L. Cao et al., Design of protein-binding proteins from the target structure alone. Nature 605, 551–560 (2022).

7. K. A. Barlow et al., Flex ddG: Rosetta Ensemble-Based Estimation of Changes in Protein-Protein Binding Affinity upon Mutation. J Phys Chem B 122, 5389–5399 (2018).

